# Effects of forest urbanization on the interplay between small mammal communities and their gut microbiota

**DOI:** 10.1101/2023.09.07.556680

**Authors:** Marie Bouilloud, Maxime Galan, Julien Pradel, Anne Loiseau, Julien Ferrero, Romain Gallet, Benjamin Roche, Nathalie Charbonnel

## Abstract

Urbanization significantly impacts wild populations, favoring urban dweller species over those that are unable to adapt to rapid and abrupt changes. One possible explanation for differential adaptative abilities between these species is that the microbiome may modulate the host phenotype rapidly through a high degree of flexibility. Conversely, under such anthropic perturbations, the microbiota composition of some species could be disrupted, resulting in dysbiosis and negative impacts on host fitness, potentially causing local extirpation. The links between the impact of urbanization on host communities and their gut microbiota have only been scarcely explored. In this study, we tested the hypothesis that the gut microbiota (GM) could play a role in host adaptation to urban environments. We addressed this question by studying several species of small terrestrial mammals sampled in forested areas along a forested gradient of urbanization (from rural forests to urban parks) during 2020 fall. The gut was collected and bacteria were described using a 16S metabarcoding approach. We tested whether urbanization led to changes in small mammal communities and in their GM. We analyzed these changes in terms of the presence and abundance of taxa and their putative functions to decipher the processes underlying these changes. We found that urbanization had marked impacts on small mammal communities and their GM, either directly or indirectly depending on small mammal species categories. The urban dweller species had a lower taxonomic diversity but a higher functional diversity and a different composition compared to urban adapter species. Their GM assembly was mostly governed by stochastic effects, which could indicate dysbiosis in these urban species. Selection processes and an overabundance of functions were detected that could be associated with adaptation to urban environments despite potential dysbiosis. Urbanization could also impact the diversity and taxonomic composition of GM in urban adapter species. However, their functional diversity and composition remained relatively stable. This observation can be explained by functional redundancy, where certain taxa express the same function. This could explain the adaptation of urban adapter species in various environments, including urban settings. We can therefore assume that there are feedback loops between the gut microbiota and the host species within communities, enabling rapid and flexible adaptation.

## 1. Introduction

Urbanization, the process of making an area more urban through higher human population within cities (Verrelli et al., 2022), is a major driver of global land use change. It is associated with increased rates of habitat loss or fragmentation, often coupled with diversity loss and species extinction (e.g. Czech et al., 2000; Dri et al., 2021; Moll et al., 2019). Global assessments report that urban expansion can be responsible for a 50% loss of local species richness (Li et al., 2022). Urbanization may induce rapid and abrupt changes that hinder the adaptation of species reliant on specific natural environments (Parmesan, 2006), ultimately leading to the extinction of entire populations (Sol et al., 2017). Species that are highly sensitive to urbanization-related changes and are unable to survive in urban areas are called ’urban avoiders’ (Fischer et al., 2015). On the opposite, some species benefit from these anthropogenic impacts. Cities can be a refuge with abundant human food and a release from biotic pressures such as predation and competition (Lowry et al., 2013; Werner & Nunn, 2020). Certain species have successfully adapted and flourished in urban environments. They attain high population densities in cities (Šálek et al., 2015; Tucker et al., 2021). When they specialize to the point of living at the expense of humans, they are named urban dwellers (Fischer et al., 2015). Species with the ability to adapt to a wide range of environments and resources (Slatyer et al., 2013) are named urban adapters (Fischer et al., 2015). These species tolerate urban conditions, in particular green areas, but also survive in rural environments.

Understanding why and how some species adapt to urbanization is pivotal to addressing issues related to biodiversity crises (Verrelli et al., 2022). Urbanization can trigger changes within species, driven by processes such as selection, epigenetic inheritance, and phenotypic plasticity (Lambert et al., 2021; Rivkin et al., 2019). Recently, Alberdi et al. (2016) advocated for a critical role of the microbial community in promoting host adaptation to rapid environmental changes, especially through its impact on host phenotypic plasticity. On one hand, the gut microbiota (noted “GM” hereafter) impacts animal biology and evolution by providing or influencing essential services that contribute to its health (Shapira, 2016) including nutrition and metabolism (Rowland et al., 2017), immunity (Belkaid & Hand, 2014) or behavior (Ezenwa et al., 2012). On the other hand, the GM is shaped by host characteristics (e.g. genomics, age or sex, Bonder et al., 2016; Santoro et al., 2017), environmental features (e.g. climatic factors, resources, Goertz et al., 2019; West et al., 2019) and their interactions (Walter & Ley, 2011). In addition, the GM can respond rapidly to environmental changes thanks to due to microbial flexibility (*i.e.* the potential for dynamic restructuring, Voolstra & Ziegler, 2020) but also to bacterial short generation time, high mutation rate.

Urbanization may induce different types of GM changes. Neutral changes are observed when the initial GM community is constant from a taxonomic or functional point of view (Moya & Ferrer, 2016b). Such changes could increase hosts’ phenotypic plasticity, which may be a prerequisite for adaptation (Alberdi et al., 2016).

Adaptive changes may occur, modifying the composition and diversity of GM. They can allow hosts to survive in new ecological niches, notably through the ability to digest new food sources, to improve metabolic capacities or to increase tolerance to deleterious environmental conditions (Moeller & Sanders, 2020). Over several generations, selection may favor hosts with advantageous GM, especially in disrupted environments (Alberdi et al., 2016; Moya & Ferrer, 2016a). In the long term, there may be congruence between the evolutionary history of various host species and the community structure of their associated microbiomes, what is named phylosymbiosis (Brooks et al., 2016; Kohl, 2020).

Lastly, maladaptive changes may also occur. They are associated with the loss of certain essential functions and the disruption of GM homeostasis. This state is named dysbiosis and has negative impacts on host fitness and health. In particular, several studies have shown that anthropogenic disturbances could lead to a decrease in GM diversity and to higher heterogeneity of microbial composition between hosts (Fackelmann, et al., 2021; Lavrinienko et al., 2021; Wilkins et al., 2019). The Anna Karenina principle (AKP) has been proposed to explain such patterns: all healthy, balanced microbiota are similar, while disrupted microbiota are all different (Zaneveld et al., 2017). Here, dysbiosis reflects stochastic processes, *i.e.,* random changes in the establishment or extinction of bacterial taxa. This can lead to immune responses and metabolism dysregulation (Nyangale et al., 2012), consequently impacting negatively the hosts’ health (Boulangé et al., 2016).

Predicting the impact of environmental disturbances, especially urbanization, on wildlife and microbiota has been the topic of several studies this last decade. However, many of these studies have focused on a single host species (e.g. Fackelmann et al., 2021; Stothart et al., 2019; Sugden et al., 2020; Teyssier et al., 2018; Phillips et al., 2018; Wasimuddin et al., 2022). The few studies that investigated GM variations within host communities led to incongruent patterns with either a stronger impact of host phylogeny over habitats on GM assembly (e.g. Knowles et al., 2019) or the opposite (Teng et al., 2022). Therefore, gathering more data on the relationships between urbanization, host community assembly and their GM remains critical in the domain of urban ecology (Verrelli et al., 2022).

Small mammals constitute a relevant model to test hypotheses regarding these relationships. They represent a large diversity of mammals, have colonized a wide array of habitats and exploit diverse foraging niches (Solari & Baker, 2007). Several rodents and insectivores occur in urban environments as urban adapters or urban dwellers. Besides, a recent meta-analysis performed by Santini et al. (2019) has revealed that high diet diversity was a main factor predicting rodent adaptation to urbanization. This suggests that the GM could be at the core of this adaptation to urban environment.

Here, we have investigated the interlinkages between urbanization, small mammal communities and their GM in forested areas. We analyzed the presence and abundance of rodents and insectivore’s species and the composition of their GM along a landscape gradient ranging from rural to urban forests. We first verified that small mammal species could be categorized as urban avoiders, adapters, or dwellers on the basis of their distribution along this urbanization gradient. Secondly, we investigated the variation in GM considering different factors, including host species, urban and rural sampling sites, and individual features. We performed community ecology analyses to examine the differences in the composition and diversity of GM i) between host species categories of urbanization response with regard to urban adaptation and ii) within host adapter species along this gradient. Next, we inferred the ecological processes (selection versus stochasticity) that could underly the variations observed, as well as the potential impacts of GM changes for the hosts (e.g. neutral, adaptative or maladaptive), following the decision tree detailed in Fig. 1. Overall, this study enabled to emphasize potential links between GM and small mammals responses to urbanization.

**Fig. 1.**
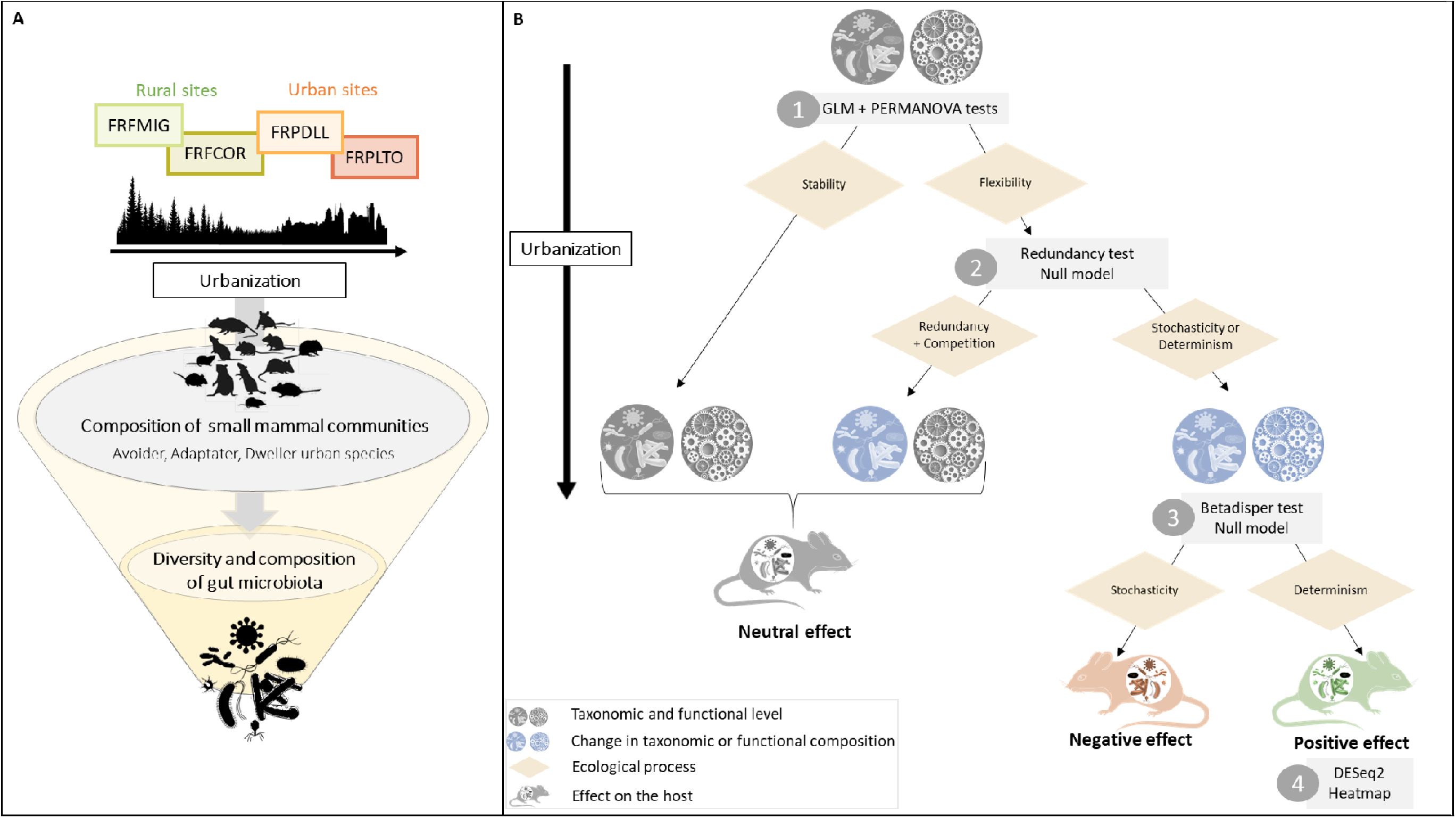
A) We are testing the impact of urbanization on the composition of small mammal communities along an 144 urbanization gradient. Additionally, we are investigating whether the diversity and composition of the gut microbiota are influenced by the sites sampled along the urbanization gradient or by the small mammal species in presence. B) Decision tree for interpreting changes in microbial composition to highlight the processes that may influence the gut microbiota and its response to urbanization. Taxonomic and functional compositions are represented by icons, with blue indicating a change and grey an absence of change. Yellow diamonds represent ecological processes, and grey rectangles indicate the statistical tests performed to infer underlying ecological processes. **Step 1.** The GLM & PERMANOVA tests assesses whether there is a change in GM diversity and composition with urbanization. If there is no taxonomic change, it suggests that the gut microbiota remains stable despite urbanization, whereas if there are changes, the GM is considered to be flexible. When taxonomic changes occur, but functional changes are not necessarily present or are weak, redundancy processes may underlie the assembly of microbial communities. **Step 2.** Redundancy analysis is applied to identify situations where taxa express the same function (considered redundant) or where each taxon expresses its own function (not redundant). This test is followed by a null model test (NTI) to determine the mechanisms that may underlie the redundancy. These mechanisms can be selective processes through overdispersion (indicating competition between taxa) or underdispersion (indicating cooperation between taxa), resulting from abiotic or biotic effects among taxa. Additionally, it is worth noting that phylogenetic dispersion between taxa can also be influenced by neutral effects. Otherwise, when taxonomic and functional changes occur, stochastic or deterministic processes may underly GM assembly. **Step 3.** The betadisper test examines differences in intragroup variances. High variance may result from strong selective pressures, such as dietary variance, or stochastic effects indicating processes favoring dysbiosis (see Anna Karenina’s principle, Zaneveld et al., 2017). Conversely, healthy species tend to express similar essential functions, resulting in lower variance. The null model helps to determine whether GM assemblage is primarily driven by stochasticity or determinism. **Step 4.** We use DESeq2 test to identify the functions that may be subject to selection. A heatmap enables to illustrate these functional changes between different species and sites along the urbanization gradient. The colored rodent icon indicates the likely impact of GM changes on its health. Grey represents a neutral effect, red a detrimental effect and green a beneficial effect.

## 2. Material and Methods

### 2.1. Data collection

#### Sampling and characterization of small mammals

We trapped small terrestrial mammals in two forests (FRFCOR: Cormaranche en Bugey; FRFMIG: Mignovillard) and two urban parks (FRPLTO: Lyon, Parc de la Tête d’Or; FRPDLL: Marcy l’étoile, Domaine Lacroix Laval) in Eastern France, in autumn 2020. All information concerning the sampling design, capture results, and individual information were detailed in Pradel et al. (2022). Sexual maturity was determined *a posteriori* based on morphological and sexual characters. Animal trapping and sample collection were conducted according the EU Directive 2010/63/EU for animal experiments, as described in Pradel et al. (2022).

The trapping success, often utilized as an indicator of relative rodent abundance when employing lethal traps, was estimated by analyzing the capture results obtained from the initial three nights of trapping (Pradel et al., 2022). It was calculated following Aplin et al. (2003) as ln(1-number of rodent trapped /(number of traps × number of nights))x(-100) (Supplementary Table S1.1).

#### Environmental characterization of sampling sites

We used Corine Land Cover (*CORINE Land Cover,* 100m of resolution) and Qgis software (Karger et al., 2017) to extract land cover area estimates for each site. In addition, we used Corine Land Cover of forests (10m of resolution) to estimate forest fragmentation using the *Landscapemetric* package (Hesselbarth et al., 2019). These calculations were made for each site by considering a 3 km buffer around the barycenter of the traps. Lastly, we extracted bioclimatic indices from the CHELSA database (1km of resolution, https://chelsa-climate.org/, Karger et al., 2017) for each site using the coordinates of the traps’ barycenter (Supplementary Table S1.2).

#### Gut microbiota sequencing

For each individual, we extracted DNA from a 5mm piece of colon with the DNeasy 96 Blood and Tissue kit (Qiagen). We followed the manufacturer’s instructions, with the exception of the addition of a bead-beating for 5 min with 500 mg of 0.5 mm zirconia beads in a TissueLyser (Qiagen) after the proteinase K digestion, as recommended in (Chapuis et al., 2023). We amplified the V4 region of the 16S rRNA gene by PCR with the primers 16S-V4F [GTGCCAGCMGCCGCGGTAA] and 16S-V4R [GGACTACHVGGGTWTCTAATCC]) following the PCR conditions and program described in Galan et al. (2016). Various controls were included to facilitate bioinformatics sorting of the sequences, including replication of libraries for all samples as well as negative controls for extraction, PCR and indexing, and the ZymoBIOMICS Microbial Community Standard (Zymo). We performed a run of 2 x 251 bp MiSeq paired-end sequencing. Information about the sequencing of samples and the raw sequence reads are detailed in the Zenodo repository (https://zenodo.org/record/8143272).

#### Sequence processing

Amplicon sequence variants (ASVs) were generated using dada2 analysis pipeline (Qiime2_2021.11) (Bolyen et al., 2019; Callahan et al., 2016). Both R1 and R2 reads were trimmed at 180 and 120 base-long respectively. This procedure improved the average quality of the reads (>Q30) and maximized the proportion of R1-R2 merging (approximately 80% of the total number of reads). Chimeric sequences were identified by the consensus method of the *removeBimeraDenovo* function. Taxonomic assignments were performed using blast+ (Camacho et al 2009) implemented in the FROGS workflow (Escudié et al., 2018) and the SILVA rRNA 138.1 database excluding the sequences with a pintail quality <100 (http://www.arb175silva.de/projects/ssu-ref-nr/). The phylogenetic tree of ASVs was built with FROGS using *FastTree* (Price et al., 2009) after a multiple alignment performed with *Mafft* (Katoh et al., 2002).

Further analyses were implemented in R v4.0.3 (R Core Team, 2020). Scripts are available in Zenodo repository (https://zenodo.org/record/8143272). Sample metadata, abundance table, taxonomy table and tree are linked in the phyloseq object using the *Phyloseq* package (McMurdie & Holmes, 2013).

We filtered false positives following the strategy described in Galan et al. (2016). In short, we discarded positive results with sequence counts below two ASV-specific thresholds, which checked respectively for (1) cross-contamination between samples or the presence of kitome during DNA extraction and PCR steps (T_CC_ for Threshold cross-contamination), and (2) incorrect assignment due to the generation of mixed clusters on the flowcell (Kircher et al., 2012) during Illumina sequencing (T_FA_ for Threshold false-assignment). The T_CC_ corresponds to the maximum number of reads observed in a negative control for DNA extraction or negative control for PCR for each ASV. The T_FA_ corresponds to the putative maximum number of reads assigned by mistake to a wrong sample. We have chosen a *Mycoplasma capricolum* sample as an internal DNA control, with a maximum rate of read misassignment of 10^-4^ in all MiSeq sequencing runs conducted in our laboratory from 2020 to 2022. Finally, for each sample and each ASV, only the occurrences confirmed by the two technical replicates were kept in the dataset. At this stage, the reads of the technical replicates for each sample were summed.

Based on the taxonomy of ASVs, only the kingdom Bacteria was retained. The chloroplast phyla, the unaffiliated phyla and the *Mitochondria* family were removed. We filtered individuals based on the rarefaction procedure implemented in *Phyloseq*. Individuals for which the sequencing depth was insufficient to determine all the ASVs present (*i.e.* for which the plateau had not been reached) were deleted. Next, we removed individuals corresponding to a low number of captures per site and species (threshold of five individuals/species/sites). We also excluded individuals found dead in the traps. Their GM composition differed from that of individuals captured alive, what could be explained by an advanced stage of degradation of the GM of these former individuals (Li et al., 2020). We also removed *Glis glis* individuals from the statistical analyses dedicated to GM, as the GM of this species exhibited an extremely low level of diversity with an overrepresentation of a particular ASV (Supplementary File 1).

ASVs number of reads were finally normalized to proportional abundance for each individual (McKnight et al., 2019).

### 2.2. Statistical analyses

#### Composition of small mammal communities

We primarily tested whether the sampling sites were representative of an urbanization gradient rather than of other potential confounding factors. After removing significant covariates detected by a Pearson correlation test, we performed a principal component analysis (PCA) with all environmental and bioclimatic features using *FactoMineR* package (Lê et al., 2008). We used the coordinates of the first axis as a score of urbanization.

Next, we analyzed whether small mammal communities’ composition was influenced by urbanization, and we verified that host species could be classified into three classical categories of urbanization response, namely avoiders, adapters and dwellers (Fischer et al., 2015). We utilized the Bray-Curtis index to generate a dissimilarity matrix of small mammals trapping success (abundance proxy) among sites. We tested whether the urbanization score influenced this dissimilarity matrix using the *capscale* function of the *vegan* package (Oksanen et al., 2020). Significance was assessed using 10,000 permutations. We selected the best model using the *ordiR²step* function.

#### Gut microbiota taxonomic and functional diversity

Throughout this study, we described GM using information relative to bacterial taxa (*i.e.* ASVs) and functional metagenomic predictions obtained using *Picrust2* software (Douglas et al., 2018). For functional predictions, the metabolic pathway description and enzyme classes were obtained from the MetaCyc database (Caspi et al., 2014).

#### Variations in the alpha diversity of the gut microbiota

We first estimated the GM alpha diversity, *i.e.* the diversity of within-host bacteria, using the taxonomic richness, the Shannon index and two metrics considering taxa phylogenetic diversity, estimated using the *picante* package (Kembel et al., 2010). These metrics are the Faith index which corresponds to phylogenetic diversity (PD) and the Nearest Taxon Index (NTI) (Webb, 2000), which reflects phylogenetic structuring near the ends of the tree. Finally, in order to address functional diversity, we calculated the richness index of predicted functions using *MiscMetabar* (e. g. Alberdi & Gilbert, 2019).

We next tested the relative influence of host species, sampling sites, and their interaction on the taxonomic and functional diversity indices described above using generalized linear models (GLMs). Individual factors such as maturity and sex of animals were also investigated. We used the negative binomial distribution for species richness and the Gaussian distribution for the other indices. We highlighted the deviation and dispersion of the model residuals from normality with the *DHARMa* package (Hartig, 2022) and we applied transformation when necessary. The selection of the best model was made by considering all possible model combinations using the *dredge* function of the *MuMIn* package and the Akaike information criterion corrected for small sample size AICc (Johnson & Omland, 2004). We then averaged the partitioned variance of each factor in the best model using *variancePartition* (Hoffman & Schadt, 2016). When site and/or species factors were significant, Tukey’s post-hoc tests were applied to assess pairwise differences in modalities, using the *multcomp* package (Hothorn et al., 2008). Because species were not evenly distributed among sites, the dataset was not balanced. To better assess the interaction effect, a factor combining species and site was defined (named species-site hereafter). A linear model was applied, followed by a Tukey post-hoc test comparing only possible biological interactions with a contrast matrix. Specifically, we compared the diversity i) between host species and ii) within adapter species along the urbanization gradient.

#### Ecological processes shaping the diversity of the gut microbiota

We used two approaches to infer the relative influence of ecological processes in shaping the GM taxonomic and functional diversity.

We examined the importance of functional redundancy by analyzing the correlations between functional and taxonomic richness for each species-site combination (Fig. 1B, step 2). A slope lower than 1 indicated functional redundancy, *i.e.* multiple bacterial taxa having the same function (Lamothe et al., 2018), while a slope greater than 1 indicated that bacterial taxa may have more than one function. We tested whether these slopes differed significantly between species-site combinations, using an analysis of covariance (ANCOVA) performed with a GLM. A post-hoc test was applied using *emmeans_test* with *rstatix* package (Kassambara, 2022) to compare pairs of slopes between species-site combinations. Here, we considered the taxonomic richness as a covariate of functional richness. The matrix contrast of biological pairs and Benjamini-Hochberg corrections for multiple tests were implemented to assess the significance of interactions.

Next, NTI values provided information about the relative influences of stochastic and deterministic processes on GM composition (Webb et al., 2002). When GM is predominantly influenced by stochastic processes, its phylogenetic composition is expected to align closely with random community assembly expectations (-2 < NTI < 2). In the opposite, selection mediated by environment (environmental filtering) should lead to phylogenetic clustering, *i.e.* the coexistence of taxa that are more closely related than expected by chance (NTI > 2). Selection mediated by bacterial species competitive interactions should lead to phylogenetic overdispersion, *i.e.* the coexistence of taxa that are more distantly related than expected by chance (NTI < -2). We tested these hypotheses by comparing ASVs phylogeny to a null model, generated by randomizing the bacterial ASV labels between taxa (with the parameter: null.model="taxa.labels"; Fig. 1B, step3).

#### Gut microbiota taxonomic and functional composition

##### Variations in the beta diversity of the gut microbiota

We measured GM beta diversity, which refers to bacteria diversity between hosts, to compare taxonomic and functional composition with regard to host species, sampling sites and individual features. Additionally, we examined i) between host species categories of urbanization response and ii) within host adapter species along the urbanization gradient.

Dissimilarities in taxon composition (ASVs) were calculated based on the normalized ASVs abundance table and the weighted unifrac index. This index considers both the abundance of ASVs and their phylogenetic relationships. We analyzed changes in GM composition along the urbanization gradient using the *vegan* package (Oksanen et al., 2020). We tested the influence of small mammal species, sites and individual factors (sex and maturity) on GM composition using a permutational analysis of variance (PERMANOVA) implemented with the *capscale* function. The best model was selected using permutation tests in constrained ordination, and the *ordiR²step* function maximizing the *R*² value. Besides, we performed a distance-based redundancy analyses (db-RDA) from the previously obtained constrained ordination. This enabled to highlight, in addition to the “*pairwise adonis*” post-hoc test, the variation between each pair within sites and species.

##### Ecological processes shaping the composition of gut microbiota

We applied several approaches to infer the relative influence of ecological processes in shaping the composition of the GM (Fig. 1B).

First, PERMANOVA analyses were performed to provide information on GM flexibility (Fig. 1B, step 1). We considered both variations in GM composition along the urbanization gradient for adaptive species and variations in GM composition between sympatric urban adapter and dweller species.

Second, we tested whether the variations in GM composition resulted from adaptive or non-adaptive changes. We estimated intra-host species and intra-site dispersal using the *PERMDISP2* test implemented with the *betadisper* function. High dispersal indicated that the GM composition was likely to be driven by stochastic processes while low dispersal could reflect selective processes favoring similar microbial functions (Fig. 1B, step 3).

In addition, the null model approach was applied to quantify the contribution of ecological processes (stochasticity vs selection) on GM composition assembly and turnover (Stegen et al., 2013). We followed the procedure described by Barnett et al. (2020) to generate the β- nearest taxon index (βNTI). Briefly, we used the *comdistnt* function from the *picante* package to determine the β-mean-nearest taxon distance (βMNTD). We generated null values of βMNTD by randomly reshuffling 1,000 times the extremities of the phylogenetic tree. Finally, βNTI was calculated according to the formula: βNTI= (βMNTDobserved-mean (βMNTD null))/ standard deviation (βMNTDnull). Selection was a major process when βNTI was lower than - 2 or higher than 2 (Stegen et al., 2013). In this case, the phylogenetic turnover of the GM was lower or higher than expected by chance respectively. In contrast, when βNTI values ranged between - 2 and 2, we concluded that stochastic processes were the main drivers of the GM composition.

After clustering individuals by species-site combination, we counted the number of observations of each ecological process, according to the βNTI values, within each combination. We performed a chi-2 test to detect observations that were significantly different from null expectations.

Lastly, we determined whether individuals from a given species-site combination had more similar functions than individuals from different species-site combinations, which could be the result of selective processes (Fig. 1B, step 4). This analysis was performed using the *DESEq2* package with count data and a negative binomial family (the Wald Test parameter) (Love et al., 2014). We added +1 to the abundance dataset to avoid a zero-inflation bias.

## 3. Results

### 3.1. Urbanization influences small mammal communities

Based on a sampling effort of 2163 traps-nights, we recorded a total of 228 small mammals*, i.e.* a global trapping success rate of 10.5%. They corresponded to 14 species. Nine species belonged to the order *Rodentia* and to four families: *Gliridae* (*Glis glis*), *Sciuridae* (*Sciurus vulgaris*), *Muridae* (*Rattus norvegicus, Mus musculus*, *Apodemus sylvaticus* and *Apodemus flavicollis*) and *Cricetidae* (*Myodes glareolus, syn. Clethrionomys glareolus*, *Microtus arvalis* and *Microtus agrestis*). Five species belonged to the order *Soricomopha* and *Soricidae* family (*Sorex araneus*, *Sorex coronatus*, *Neomys fodiens*, *Crocidura russula* and *Crocidura leucodon*) (Fig. S1).

#### Sampling sites delineate an urbanization gradient

The first axis of the PCA based on site characteristics explained 60.97% of the total variation and opposed artificial areas and contiguous forests to a forested environment with larger forested areas (Fig. 2A). Sampling sites were organized along this axis that described a gradient of urbanization with urban sites (FRPLTO and in a lesser extent FRPDLL) and rural sites (FRFCOR and FRFMIG).

**Fig. 2.**
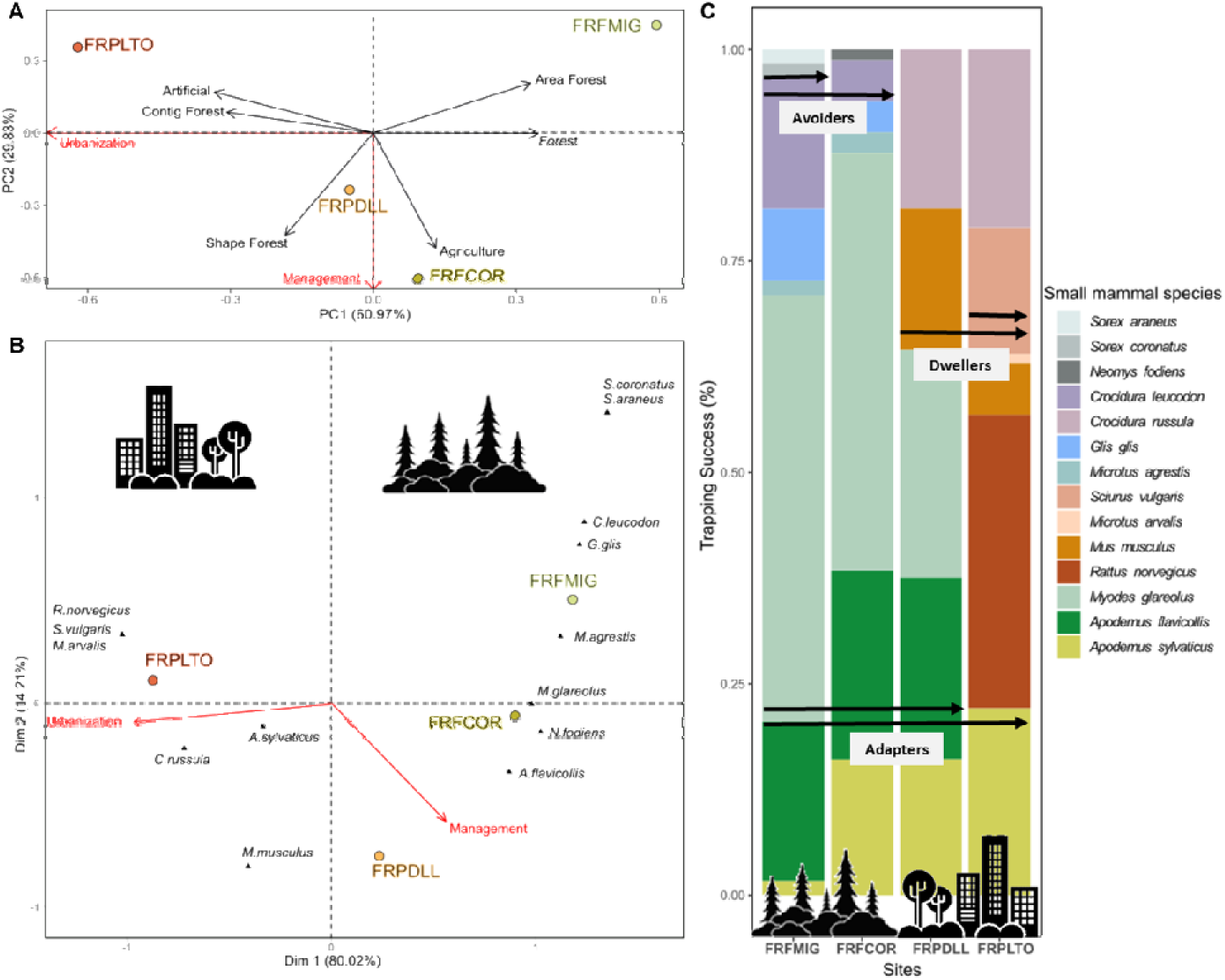
Analyses of small mammal community variations. A) Principal component analysis (PCA) of the environmental features characterizing sampling sites (black arrows. The red arrows indicate that PCA1 axis describes a gradient of urbanization while PCA2 axis describes differences in land management. B) Canonical correspondence analysis (CCA) of the small mammal trapping success per site. Sites are represented by dots and color code. Species are represented by a black triangle. The two first PCA axes were included as explanatory variables in the canonical analysis and are represented by red arrows. C) Bar graph showing the relative trapping success of species (represented by colors) per site, ordered from the less (FRFMIG) to the more (FRPLTO) urbanized ones. The icons highlight the variations between urban and rural locations, with arrows denoting the species classification on the urbanization spectrum. Species that are not found in urban zones are classified as urban avoiders, those only present in urban areas are urban dwellers, and those found in both areas are urban adapters.

#### Small mammal species exhibit distinct categories of response to urbanization

The assembly of small mammal communities differed among sampling sites (Fig. 2A; Fig. S2). Although the geography, climate and land use explained part of this variation (Fig. S3; Supplementary Table S1.2), urbanization was the main factor driving the differences observed between sites. Rural sites homed seven (FRFCOR) or eight (FRFMIG) small mammal species and they had six species in common. Urban sites homed fewer small mammal species (respectively five in FRPDLL and six in FRFLTO) and they shared three species. FRFDLL also had three species in common with the rural sites, while FRFPLTO had only one species in common with these sites.

The CCA based on small mammal relative abundance revealed that sites differed in the composition of small mammal communities and this variation was found to be associated with urbanization (Fig. 2B). The first axis of the CCA explained 80% of the total variance and it represented the urbanization score. PERMANOVA tests showed that small mammal community composition was significantly influenced by urbanization (F=5.36, p=0.04) with 57% of the variance explained by this factor (Supplementary Table S1.3).

Small mammal species corresponded to the avoider, adapter and dweller categories defined in the literature to describe wildlife responses to urbanization (Fischer et al., 2015). Some species, *G. glis*, *M. agrestis*, *S. araneus*, *S. coronatus*, *N. fodiens* and *C. leucodon,* were urban avoiders. Their relative abundance was always lower than the urban dweller and adapter species. Other species were present in both rural and urban sites. These urban adapters are *M. glareolus*, *A. flavicollis* and *A. sylvaticus* (Fig. 2C). They varied in abundance between sites: the relative abundance of *A. sylvaticus* increased along the urbanization gradient while the relative abundance of *M. glareolus* and *A. flavicollis* decreased with urbanization. They were even absent from the most urbanized site FRPLTO. In contrast, urban dwellers, namely *R. norvegicus*, *M. musculus*, *S. vulgaris* and *C. russula,* were relatively abundant compared to urban adapters.

Rare species were not included in further analyses of GM diversity and composition.

### 3.2. Urbanization influences alpha diversity of small mammal gut microbiota

We obtained a total of 5478 ASVs derived from 222 small mammals after the filtration steps. They corresponded to 1969 enzyme commissions (ECs) and 383 metabolic pathways. For all analyses, we found similar results between EC and metabolic pathway diversity, so we only presented the results relative to pathways below.

#### Small mammal species identity influences gut microbiota alpha diversity (Fig.1B, step1)

Small mammal species was the main factor explaining variations in GM alpha diversity, whatever the indices analyzed (GLM, taxonomic richness: *F*=80.42, *p=*<2.2e-16; Shannon index: *F*=56.75, *p=*2.2e-16; Faith’s PD: *F*=37.58, *p=*2.2e-16, Fig. 3A; functional diversity: *F*=7.29, *p=*4.4e-07; Fig. 3B; Table1).

**Fig. 3.**
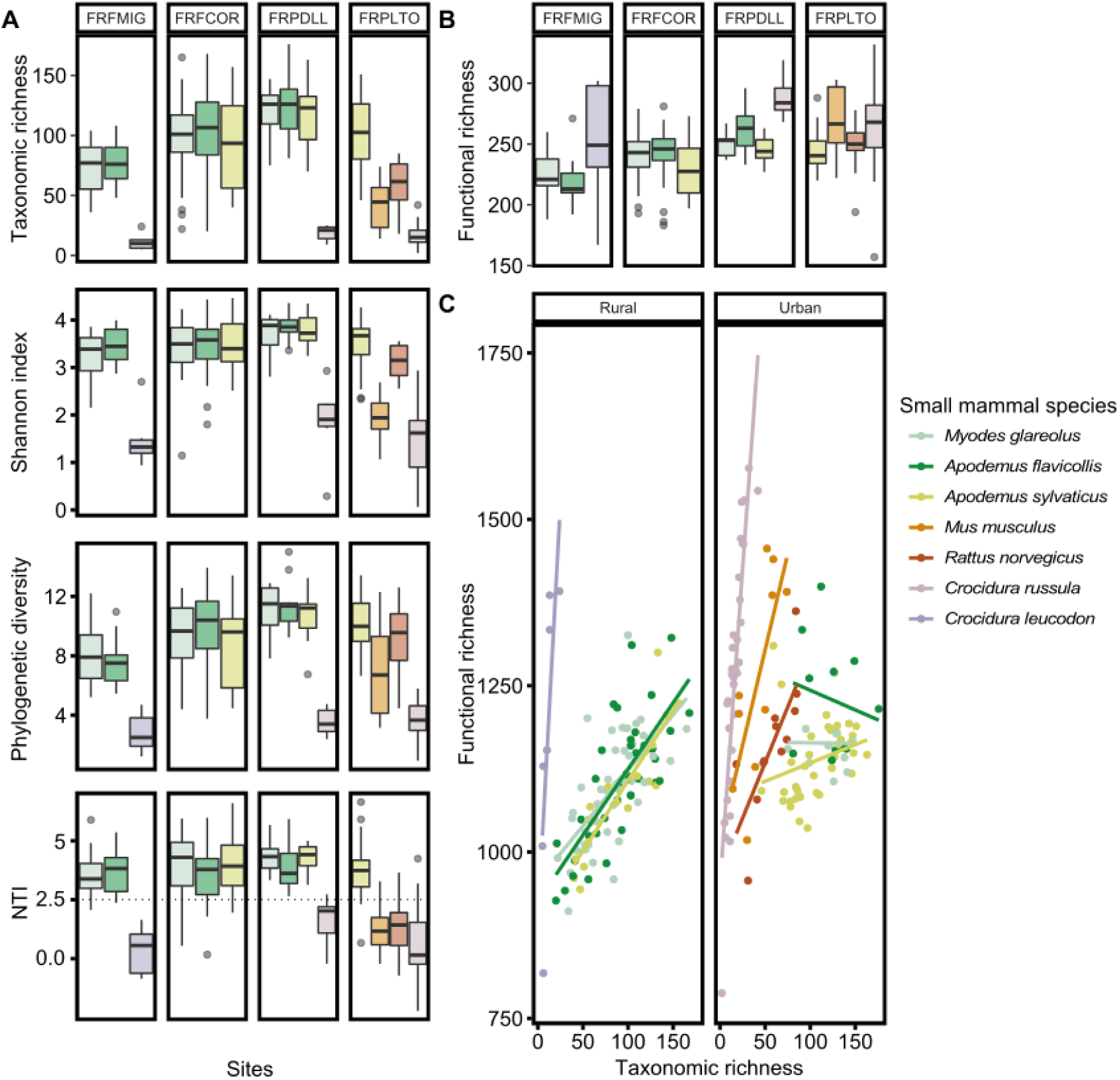
Boxplot showing variation in the alpha diversity of the gut microbiota between small mammal species in rural sites (sites FRFMIG and FRFCOR) and urban sites (sites FRPDLL and FRFLTO). The color codes correspond to small mammal species and they remain the same for all subfigures. Alpha diversity was measured A) at the taxon level with the taxonomic richness (number of ASVs), Shannon index, phylogenetic diversity (PD) and Nearest Taxon Index (NTI) and B) at the functional level with the functional richness corresponding to the number of metabolic pathways. C) Plot of the correlation between the taxonomic and functional richness for each small mammal species in rural (FRFMIG, FRFCOR) and urban (FRPDLL, FRPLTO) sites.

**Table 1.**
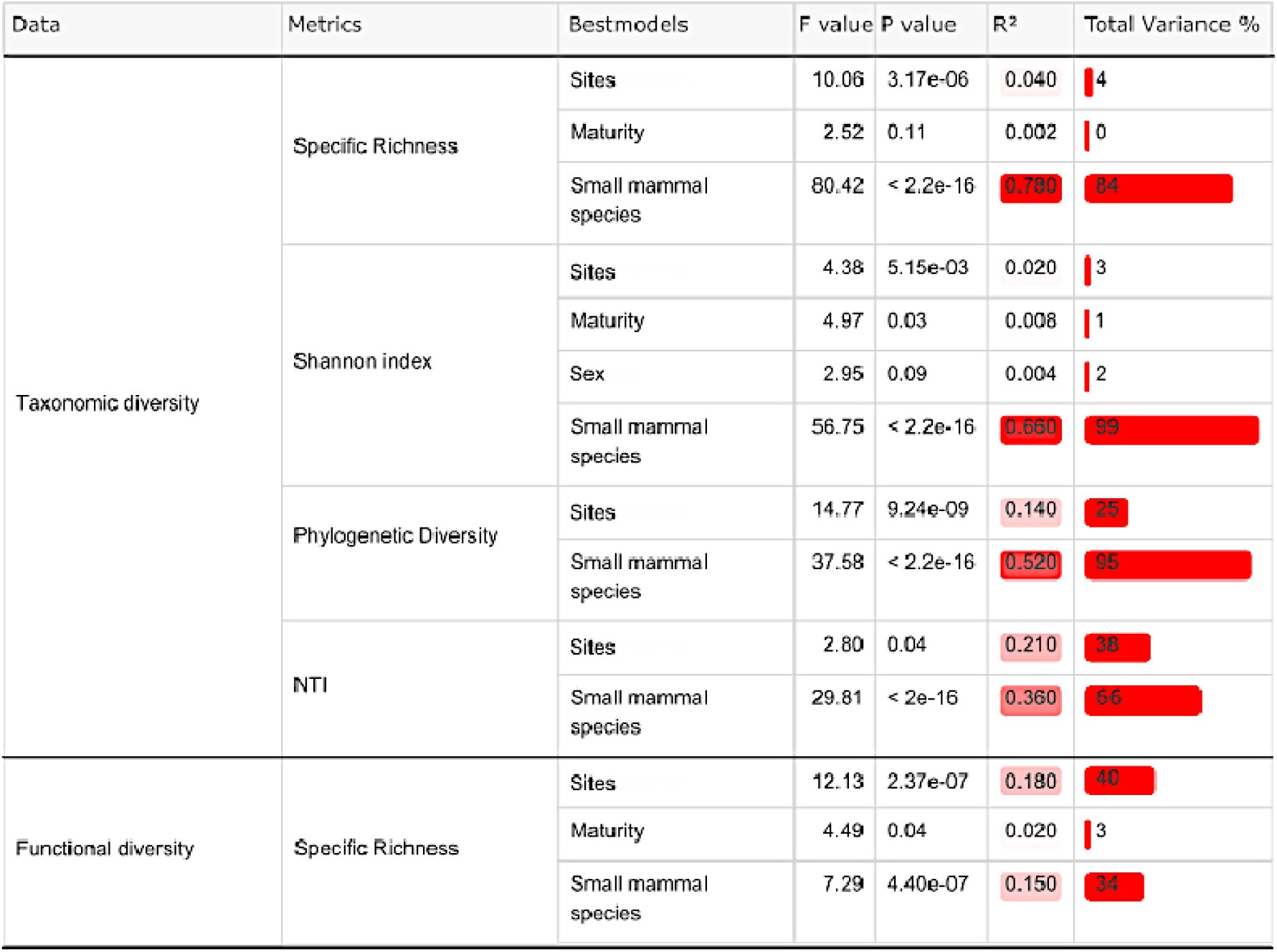
Results of the best models for the different GLM tests performed for each taxonomic and functional diversity index. For each variable in the best model, the *F* value, the *P* value, R² and the percentage of the variance explained are indicated (some factors explain the same proportion of the variance). The red gradient indicates the importance of the R² and of the variance explained in the models.

Among *Muridae*, the GM of the three urban adapter species, *A. sylvaticus*, *A. flavicollis*, and *M. glareolus*, exhibited similar levels of alpha diversity, whatever the indices considered (Fig. 3; Supplementary Table S2.1).

The GM taxonomic diversity was significantly lower for urban dweller species than for urban adapter species (Fig. 3A; Supplementary Table S2.1). *C. russula, M. musculus and R. norvegicus* had lower taxonomic diversity than *Apodemus* sp. and *M. glareolus* in FRPDLL and/or FRPLTO. This pattern was less pronounced for *R. norvegicus* in FRPLTO when considering Shannon and PD indices compared to the taxonomic richness (Fig. 3A, Supplementary Table S2.2).

Conversely, the urban dweller species showed a greater GM functional diversity than the urban adapter species (e.g. *M. musculus*, *R. norvegicus* and *C. russula* compared to *A. sylvaticus* in FRPLTO, or *C. russula* compared to *Apodemus* sp. and *M. glareolus* in FRPDLL; Fig. 3B; Supplementary Table S2.2).

#### Urbanization influences gut microbiota alpha diversity (Fig. 1B, step1)

Overall, the GM diversity increased with urbanization for all diversity indices considered except NTI (Fig. 3A and 3B, Supplementary Table S2.1). The GM diversity of adapter species was lower in the rural sites FRFMIG and FRFCOR than in the urban ones FRPLTO and FRPDLL (Supplementary Table S2.1). This impact of urbanization was dampened when considering the Shannon index (Supplementary Table S2.1).

The interaction between species and site significantly explained variations of alpha diversity in all the models tested (GLM; Taxonomic richness: *F*=43.86, *p*<2.2e-16; Shannon: *F*=33.67, *p=*2.2e-16; PD: *F*=21.39, *p*<2.2e-16; Functional richness (Pathway): *F*=9.27, *p=*5.58e-15; Supplementary Table S2.2). The GM alpha diversity of *M. glareolus* and *A. flavicollis,* but not *A. sylvaticus,* increased along the urbanization gradient whatever the indices considered (Supplementary Table S2.2).

#### Gut microbiota functional redundancy and phylogenetic clustering vary with urbanization (Fig. 1B, step2)

The covariance between functional and taxonomic richness indices was significantly influenced by small mammal species (ANCOVA, *F*=19.71, *p=*2.2e-16), site (ANCOVA, *F*=6.71, *p=*2×10^-4^) and sex (ANCOVA, *F*=5.26, *p=*0.02) (Fig. 3C). Overall, the covariance was higher for urban dweller species than for urban adapter species (Fig. 3C; Supplementary Table S2.3). This pattern was also significant at the scale of a particular site. For example, at FRPLTO, the slope for all other urban dweller species was higher than 1 (*R. norvegicus, a* =3.30; *M. musculus, a*=5.70; *C. russula, a*=17.00) whereas it was lower than 1 for *A. sylvaticus* indicating functional redundancy for this urban adapter species (*A. sylvaticus, a*=0.37).

The covariance decreased along the urbanization gradient for urban adapter species, indicating high levels of functional redundancy in urban parks but not in forests (Supplementary Table S2.3; Fig. 3C; *M. glareolus*: FRFMIG, *a*=2.10, FRFCOR, *a*=1.90 and FRPDLL, *a*=-0.02; *A. flavicollis*: FRFMIG, *a*=2.10, FRFCOR, *a*=1.90; FRPDLL, *a*=-0.59; *A. sylvaticus*: FRFCOR, *a*=2.10, FRPDLL, *a*=0.93 and FRPLTO, *a*=0.37).

NTI showed significant variations mainly among species categories of urbanization response (*F*=29.8, *p=*2.2e-16). We detected high NTI values for urban adapter species (NTI < -2), indicating a significant phylogenetic clustering (Fig. 3A. Supplementary Table S2.1). No significant pattern was detected for the urban dweller species (NTI values ranged between -2 and 2, close of 0). No changes were observed along the urbanization gradient for the adapter species (Supplementary Table S2.1).

### 3.3 Gut microbiota composition changes between small mammal species and along the urbanization gradient

#### Gut microbiota composition varies with urbanization (Fig. 1B, Step1)

The GM taxonomic composition was significantly influenced by small mammal species (PERMANOVA, *R²*=0.32, *F*=12.57, *p=*0.001) and sites (PERMANOVA, *R²*=0.08, *F*=9.06, *p=*0.001) (Supplementary Fig. S3, Table S3.1). We observed the same results for the GM functional composition (PERMANOVA, species: *R²*=0.25, *F*=8.50, *p=*0.001; sites: *R²*=0.08, *F*=7.70, *p=*0.001; Supplementary Table S3.1).

The db-RDA (Fig. 4A, 4B) showed that the clustering of individuals was driven by their species affiliation, then by their adaptation to urbanization rather than by their phylogeny (partial Mantel test, ASV: *R*=0.61, *p=*0.004; function *R*=0.58, *p=* 0.004; Supplementary Fig. S5, Table S3.2). Indeed, the first axes opposed urban dweller species to urban adapter species.

**Fig. 4.**
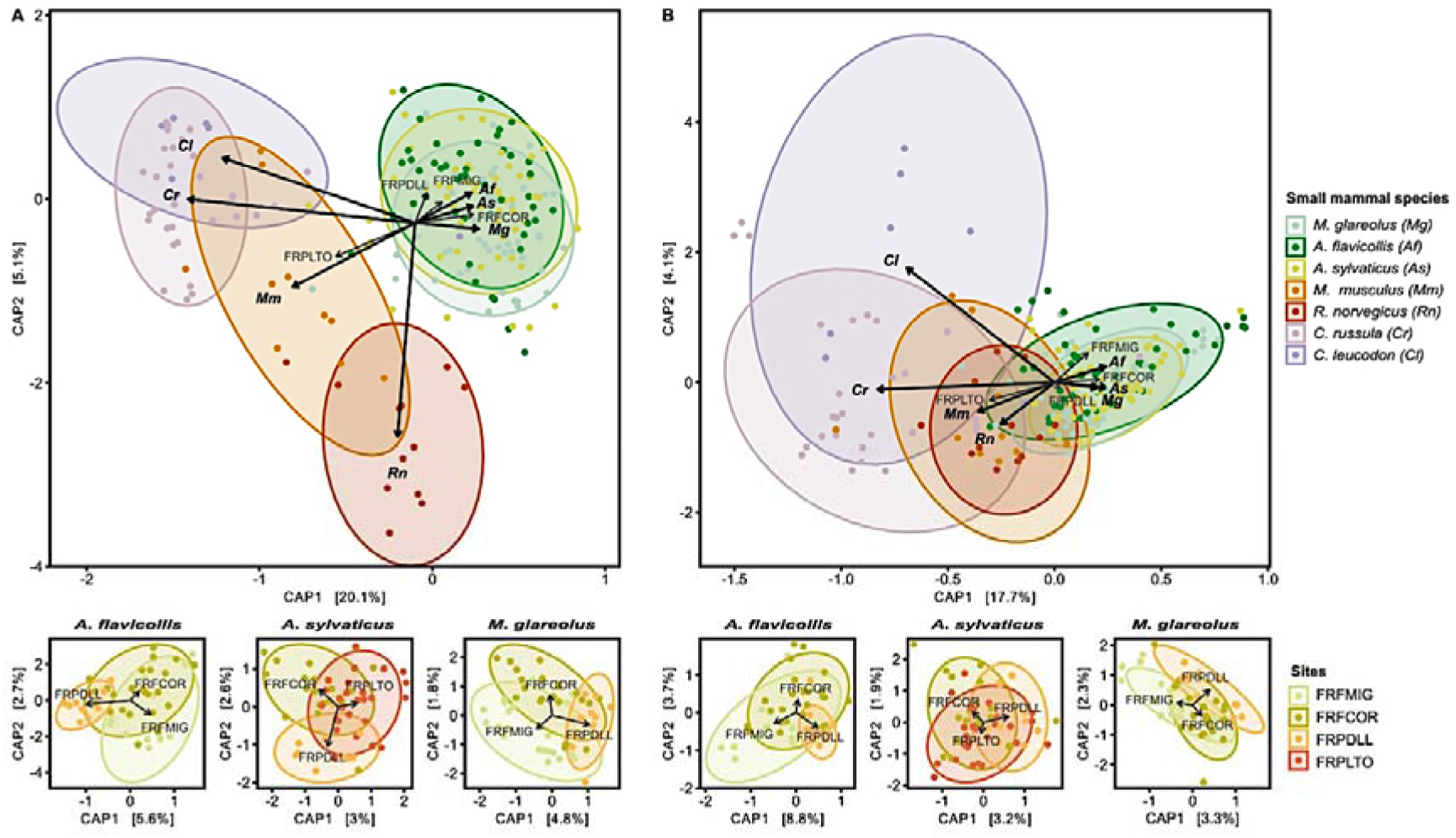
Distance-based redundancy analysis (db-RDA) of small mammal GM performed A) on ASVs and using the weighted Unifrac dissimilarity matrix and B) on functions and using the Bray-Curtis dissimilarity matrix. Only significant factors based on the *capscale* and *ordiR2step* analyses are indicated by arrows. The ellipses represent a 90% confidence interval around the centroids of the groups. The larger graph shows the significant factors modulating GM composition. Each point represents the GM of an individual and the color illustrates the small mammal species. The smaller graphs below represent the variation of the GM composition along the urbanization gradient for urban adapter species (*A. sylvaticus; A.flavicollis* and *M. glareolus*). Each point represents the GM of an individual and the colors represent sites and the urbanization gradient.

The GM composition (ASVs and functions) of *A. sylvaticus* (family *Muridae*) was closer to the one of *M. glareolus* (family *Cricetidae*) than to the one of *R. norvegicus* or *M. musculus* (family *Muridae*) (Fig. 4). These differences were also observed when considering species living in sympatry, especially at FRPLTO (Supplementary table S3.1, Fig. S4).

Urban adapter species showed contrasted changes in GM composition along the urbanization gradient. No change in taxonomic nor functional composition was observed for *A. sylvaticus* (ASV: Fig. 4A, *F*=1.60, *p=*0.052; function: Fig. 4B, *F*=1.46, *p=*0.110). *A. flavicollis* GM composition slightly changed with urbanization at both the taxonomic (Fig. 4A, *F*=2.03, *p=*0.020) and functional (Fig. 4B, *F*=3.22, *p=*0.010) levels. *M. glareolus* GM changed with urbanization at the taxonomic level (Fig. 4A; *F*=1.87, *p=*0.020) but not at the functional level (Fig. 4B; *F*=1.58, *p=*0.090).

The relative influence of ecological processes driving gut microbiota composition varies with urbanization (Fig. 1B, step3). We showed that the overall dispersion of the GM composition tended to increase with urbanization (Betadisper test, ASV: *F*=17.53, *p=*3.16e- 10; function *F*=11.73, *p=*3.78e-07, Fig. S6), in particular at FRPLTO (Supplementary Table S3.3). Small mammal species differed significantly in terms of dispersion (Betadisper test, ASV: *F*=6.67, *p=*9.37e-08; function: *F*=31.85, *p=*2.2e-16). Urban adapter species exhibiting lower levels of dispersion than some urban dwellers, mainly in urban park (e.g. *C. russula* for both taxonomic and functional composition; *M. musculus* for the functional composition only, Supplementary Table S3.3).

When we considered species-site combinations, we showed that urbanization dampened the dispersion in GM composition for two urban adapter species (Betadisper test, ASV: *F*=5.06, *p=*1.77e-06, Fig. 5A; function: *F*=16.30, *p=*2.2e-16, Fig. 5B; Supplementary Table S3.3), *M. glareolus* and *A. flavicollis* (Betadisper test, ASV: *F*=5.06, *p=*1.77e-06; function: *F*=16.30, *p=*2.2e-16, Fig. 5A.B; Supplementary Table S3.3).

**Fig. 5.**
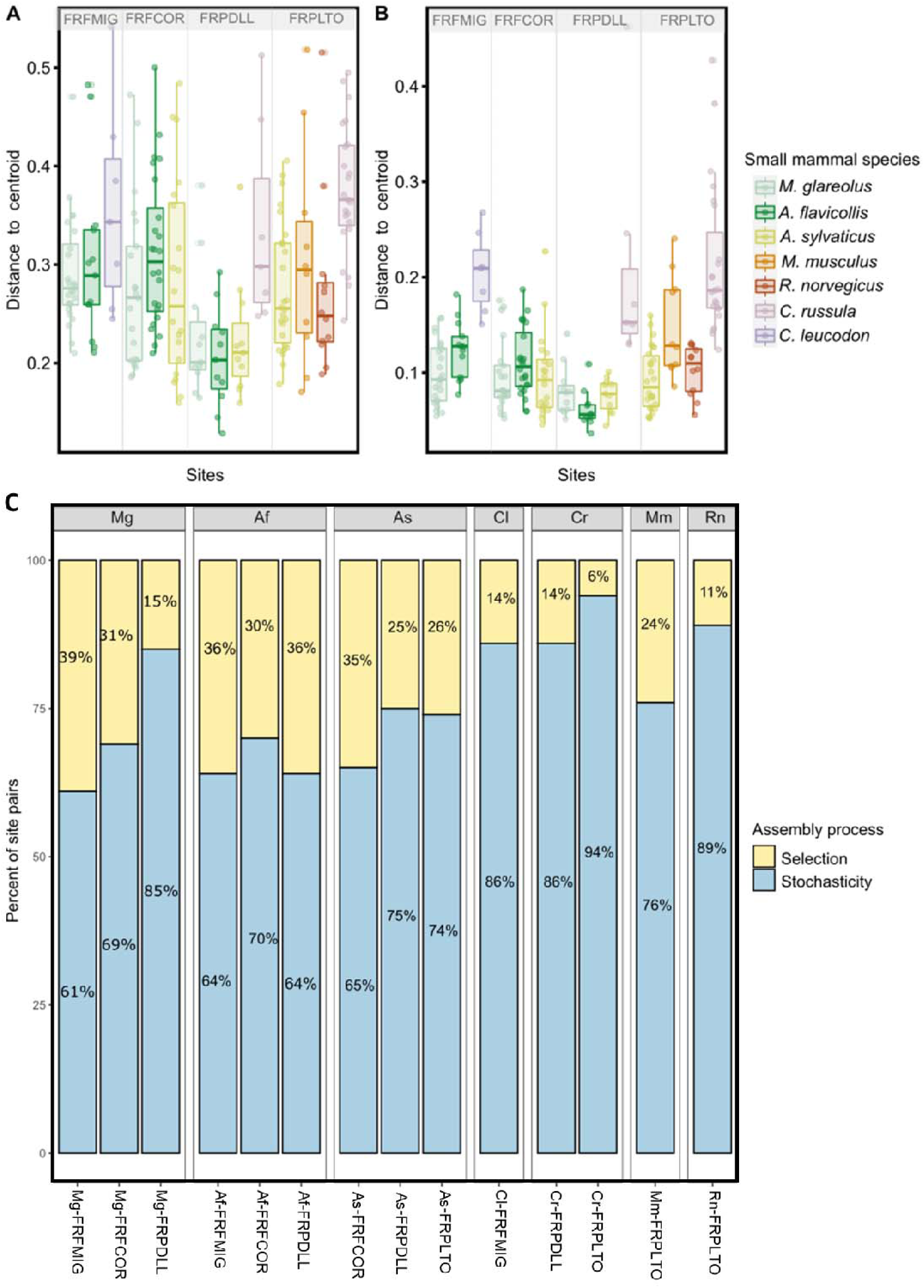
Boxplot of the distance to the centroid of each individual for each species-site combination at A) taxonomic and B) functional levels. Each point corresponds to an individual and the colors correspond to small mammal species. The combinations are ordered according to the urbanization gradient. C) Processes responsible for bacterial assembly and turnover in each combination of small mammal species and site. Percentage of pairs of individuals within each species-site combination for which selective processes (yellow) or stochastic processes (blue) are emphasized.

We next assessed the relative influence of community assembly processes using the βNTI index. Phylogenetic turnover was mainly shaped by stochasticity (>60%) (Fig. 5C). The allocation of assembly processes within species-site combinations was significantly different from what was expected (Chi-2 test, *c^2^*= 120.5, *p=* 2.2e-16). The “small mammal species” and “sampling sites” factors had significant effect on GM assembly. The GM assembly of urban adapter species was relatively more influenced by selection in forests (FRFMIG: *M. glareolus* 39%, *A. flavicollis* 36%; FRFCOR: *M. glareolus* 31%, *A. flavicollis* 30% and *A. sylvaticus* 35%), as well as in FRPDLL for *A. flavicollis* (36%) (Fig. 5C, S7). We detected a strong influence of stochasticity in dweller species (*C. russula* in FRPDLL and FRPLTO, 86% and 94%, respectively; *R. norvegicus* in FRPLTO, 89%), avoider species (*C. leucodon* in FRFMIG, 86%) and for *M. glareolus* in the urban park FRPDLL (85%) (Fig. 5C, S7).

The abundance of gut microbiota pathways and enzymes varies with urbanization (Fig. 1B, step4). We found 236 out of 383 metabolic pathways and 1064 out of 1969 enzymes with differential abundances between species-site combinations (Supplementary Table S3.4, Fig. 6; Fig. S8B). The urban dweller species diverged from urban adapter species due to an over-representation of some hydrolases (ester bond group) and transferases (sulfur and phosphorus containing groups, acyltransferases, glycotransferases). Different enzyme classes showed differential abundance along the urbanization gradient for the three urban adapter species. Hydrolases (ester bond group) were more abundant in urbanized sites for *M. glareolus* and *A. sylvaticus*. For *M. glareolus*, we also detected an over-representation of lyases (C.C. group) and an under-representation of hydrolases (peptidases and carbon nitrogen group) and oxidoreductases in FRPDLL compared to FRFMIG and FRFCOR. For A. *flavicollis*, we found an over-representation of transferases (acyltransferases and one carbon) and isomerases (intramolecular lyases) in FRPDLL compared to FRFMIG and FRFCOR. The results for the taxonomic families and pathways are summarized in supplementary materials (Supplementary Table S3.4, Fig. S8A).

**Fig. 6.**
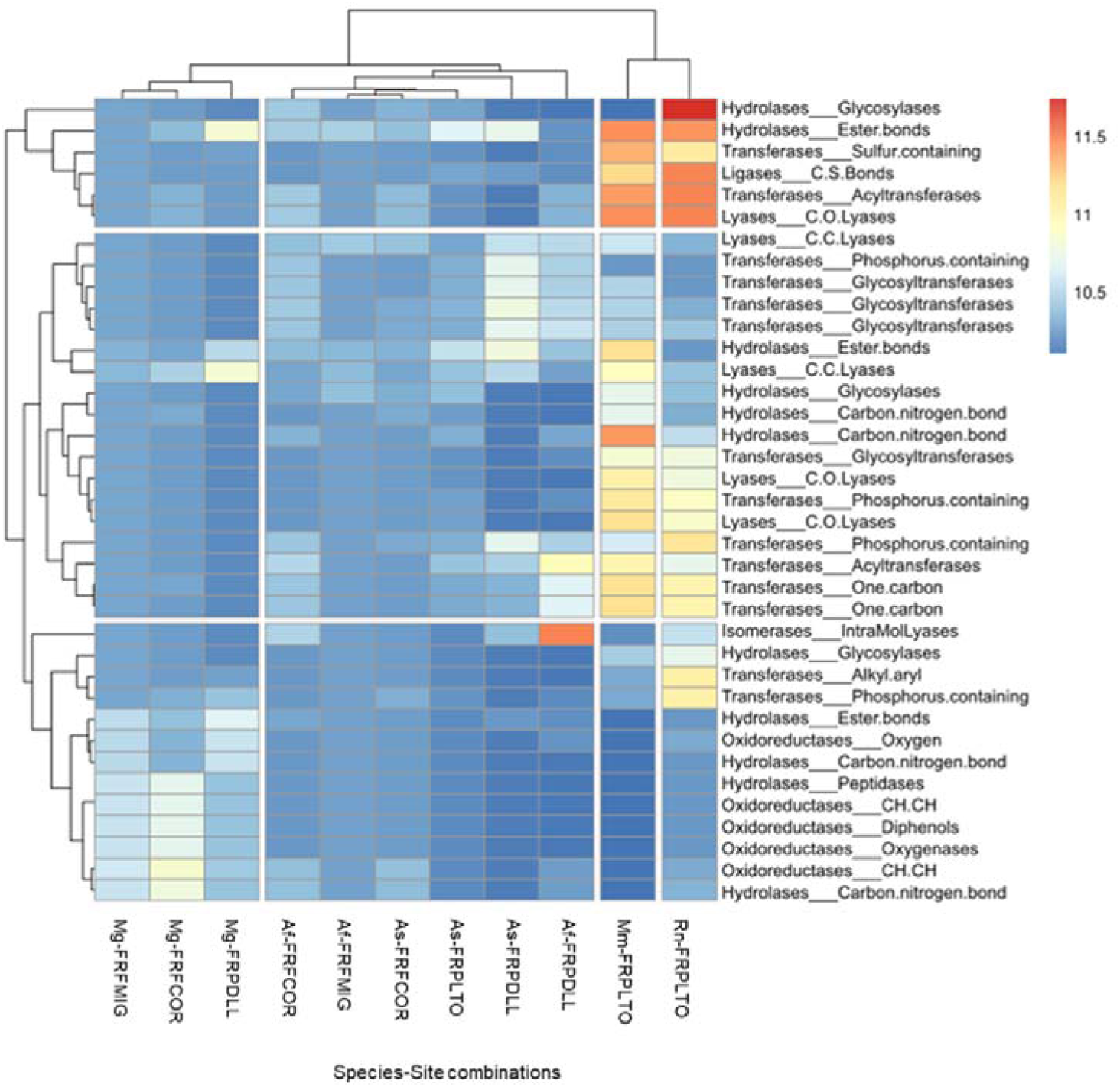
Heat map based on the DESeq2 results, considering GM functions (c-enzymes, EC) with significantly different abundances between species-site combinations. The color gradient corresponds to the magnitude of the differences. Red values indicate a strong score and blue values a weak score. Species-site combinations and enzyme classes are ordered according to their profile.

## 4. Discussion

### 4.1 Small mammal communities change with urbanization

Urbanization represents one of the most extreme forms of land use change (Johnson & Munshi-South, 2017). It may result in modifications of host community composition with an increased abundance of exotic species at the expense of native ones (Faeth et al., 2011). Here we sampled small mammals in forested sites along a gradient of urbanization, which was distinguished by the extent of artificial features as well as the size and fragmentation of the forests. We observed marked changes in the composition of small mammal communities along this gradient, as previously shown in other studies (Grimm et al., 2008; Sih et al., 2011). The presence and abundance of small mammal species along the urbanization gradient enabled to classify them into the three classically defined categories. i) The urban avoiders, *G. glis*, *M. agrestis*, *S. araneus, S. coronatus, N. fodiens*, and *C. leucodon*, which were only found in rural forests and had low abundance. ii) The urban adapters, *A. sylvaticus*, *M. glareolus*, and *A. flavicollis*, which were present in most types of forests. iii) The urban dweller species, *R. norvegicus*, *M. musculus*, and *C. russula*, which were found exclusively in cities. These categories may result from different ecology and life-history traits of small mammals, leading to the ability to adapt or not to the rapid and unfavorable changes associated with urbanization (e.g. habitat fragmentation and pollution, Grimm et al., 2008). Besides, urbanization can offer abundant and continuous resources, as well as new refuges, which can lead to the coexistence of adapter and urban dweller species (Chamberlain et al., 2009; Ofori et al., 2018). In our study, this resulted in a maximum diversity and composition of small mammal communities observed in the peri-urban park FRPDLL. This pattern has already been shown in other countries (e.g. Grade et al., 2022) and aligns with the intermediate disturbance hypothesis proposed by Connell (Sheil & Burslem, 2013).

### 4.2 Host species-driven flexibility of the gut microbiota: the influence of categories of urbanization response

Our findings showed that species identity was a foremost determinant of GM diversity and composition. As noted by Kohl (2020), the GM exhibited inter-individual variation but remains relatively conserved based on the identity of the host species. Such phylosymbiosis (observed concordance between the GM and the host species) has already been described in former studies on sympatric species of small mammals (Knowles et al., 2019; Weinstein et al., 2021).

Despite this undeniable role of host phylogeny in microbial structure, our results emphasized significant variations in the diversity and composition of the GM among different species categories, reflecting their responses to urbanization. Specifically, we observed that the structure of gut microbiota differed between categories, but were similar within each category. The urban adapter species showed a distinct microbial composition from the dweller species. *Apodemus spp* (family *Muridae*) had a microbial composition closer to *M. glareolus* (family *Cricetidae)* than to its close relatives (e.g. *R. norvegicus* and *M. musculus*, family Muridae). Varudkar & Ramakrishnan (2018) previously reported a similar pattern while comparing the GM of two phylogenetically related rats in rural and urban habitats. Habitat emerged as the main predictor of GM diversity and composition. More studies are now required to corroborate the strong impact of responses to urbanization on inter-specific differences in GM, and to determine the relative influences of deterministic and stochastic processes in shaping this microbial community in wildlife.

### 4.3 Stochasticity, a major process shaping microbial diversity and composition in urban dweller and avoider species

Our study highlighted the influence of distinct ecological processes on the GM diversity and composition of urban dweller vs adapter species. Urban dweller species had higher levels of intraspecific variance in their GM composition than adapter species. This may be due to larger ecological niche, for example with differences in diet between urban and rural areas, as has been reported in coyotes (Sugden et al., 2020). In rural forests, small mammals mainly eat seeds, fruits, and insects, whereas in urban areas, they may have access to a wider range of food sources such as garbage and pet food. However, other factors beyond diet may also be at play, as this pattern of GM variance was observed even when species were found in sympatry and potentially sharing their resources and habitats. For instance, the variance in the GM composition of wood mice was lower than that of house mice and rats, even when only considering the highly anthropized urban park FRPLTO.

Our results provided several evidence for a strong influence of stochastic processes on the GM diversity and composition of urban dweller species. Indeed, we found no strong evidence of bacterial competition or environmental filtering mediating GM composition. We demonstrated that stochasticity was more important than expected in the assembly and renewal of their GM. High variance of the GM in these species could thus reflect the deregulation of the bacterial composition and dysbiosis, with individuals acquiring different microbiota due to stochastic processes (Zaneveld et al., 2017). Although this phenomenon has already been observed in urban environments (Stothart & Newman, 2021; Sugden et al., 2020), it remains important in the future to investigate how this stochasticity and high variance in GM composition impact the physiology, health and fitness of small mammal species.

In addition, we detected higher levels of functional richness and lower levels of functional redundancy in urban dwellers compared to adapter species. Dweller species had a GM that consisted of a small number of specialized taxa with unique functions. The GM diversity and composition might hence have contributed to the specialization of these dweller species to urban environments (Moeller & Sanders, 2020). For instance, the urban dweller species showed a lower representation of Muribaculaceae and Rikenallaceae bacteria (as seen in mice fed with low fiber diet, e.g. Wang et al., 2020) and an over-representation of Helicobacteraceae, as previously found in mice fed a high-fat diet (Ali Irshad Koh Young-Sang, 2015). We also observed a higher representation of several enzymes and metabolic pathways in dwellers compared to urban adapter species. These functions could play critical role in urban dweller species adaptation in cities, in particular by counteracting the damaging anthropogenic pressures encountered in such disturbed environment. As such, in humans, genes encoding for hydrolases are overexpressed in cities, and some of the metabolic functions related to these enzymes are involved in the regulation of cardiovascular diseases, inflammatory responses and neurological diseases (Morisseau, 2022). In cities, GM may have enabled urban dwellers to settle and persist while being "sick".

The gut microbiota (GM) of the urban avoider species studied here, the soricomorph *Crocidura leucodon*, exhibited similar patterns (low alpha diversity, high inter-individual variance in GM composition) driven by the same processes (low levels of bacterial competition and functional redundancy, high rate of stochasticity) than other urban dweller species, in particular the phylogenetically close *C. russula*. This similarity may stem from ecological specialization rather than urban adaptation (e.g. Zepeda Mendoza et al., 2018). The GM may play a role in the expansion or specialization of the host niche, known as the extended phenotype (Nougué et al., 2015). In this study, *C. leucodon* was trapped in a single forest. As it habits open woodland environments and agricultural areas, it would be interesting to investigate the ability of this urban avoider species to resist or respond to rapid environmental changes, as those experienced due to climatic and anthropogenic changes.

### 4.4 Variable influence of ecological processes in shaping the gut microbiota of urban adapter species in response to urbanization

We next focused on urban adapter species and investigated GM flexibility along the urbanization gradient. We revealed an impact of urbanization on GM diversity and composition, as previously described for other mammals (e.g. Stothart et al., 2019; Sugden et al., 2020).

We observed a slight increase in GM diversity and composition with urbanization in the adapter species *Apodemus flavicollis* and *Myodes glareolus*. Higher values of GM diversity and marked differences in GM composition were observed in the peri-urban park FRPDLL compared to forests.

The patterns for taxonomic richness were less pronounced when considering the Shannon index, meaning that changes in bacterial diversity were mainly driven by the presence of rare species. These results were congruent with the observed changes in small mammal communities associated with urbanization. This could suggest a potential cascading effect of changes in host communities on host microbial communities, or a similar impact of environmental disturbance on both communities with higher species diversity maintained in environments with intermediate disturbance (Intermediate Disturbance Hypothesis, Connell, 1978; Willig & Presley, 2018). Changes in the resources available in the environment associated with anthropization could for example lead to greater microbial diversity and larger ecological niche (Weinstein et al., 2021).

These patterns were also less marked when comparing results obtained from functions and from taxonomy. Besides, we provided evidence for higher levels of functional redundancy and stochasticity in urban parks than in forests, although this result was less marked for *A. sylvaticus*. We have detected signals of phylogenetic clustering resulting from strong bacterial filtering (Webb et al., 2002). When a species of bacteria cannot fulfill a specific function due to unfavorable environmental conditions, another species or group can step in. This ensures the stability of essential functions (Konopka, 2009). However, if this functional redundancy becomes excessive, particularly due to intense environmental selection, it could lead to a homogenization of functions, making the microbiome less resilient to unexpected environmental changes. This urban homogenization has been observed in urban microbial soil (Delgado-Baquerizo et al., 2021). Lastly, the relative importance of selection in shaping the GM of these urban adapter species was slightly higher in forests than in parks, and we detected changes in enzymes and metabolic pathways along the gradient of urbanization, some of them being shared by at least two of these rodent species (e.g. PWY.5005, PWY.6628 or hydrolases ester bounds for *M. glareolus* and *A. sylvaticus*) and other being specific.

Altogether, these results indicated that the GM diversity and composition of urban adapter species are quite stable along a forest gradient of urbanization. Surprisingly, we found no evidence of dysbiosis in urban sites for these species, contrary to what could have been expected from other studies (e.g. for the spiny rat *Proechimys semispinosus,* Fackelmann et al., 2021). This suggests GM resistance or resilience (Moya & Ferrer, 2016b; Sommer et al., 2017) in the face of such environmental changes (Alberdi et al., 2016; Moeller & Sanders, 2020; Voolstra & Ziegler, 2020b), with the impact of stochasticity being dampened by functional redundancy (Moya & Ferrer, 2016b; Sommer et al., 2017). Temporal surveys would be necessary to corroborate this effect of urbanization on GM dynamics and evolution in urban adapter species, and to better understand how these responses to urbanization may affect their fitness.

## 5. Conclusions

This study corroborated the strong interlinkages existing between small mammal communities and their GM, that might change with hosts’ and abiotic features. As such, feedback loops might occur between host communities and their microbiota in response to environmental perturbations (Miller et al., 2018; Moorhead et al., 2017). In particular, urbanization seemed to impact the diversity and composition of GM differently, either as a cause or a consequence of the host species’ ability to cope with environmental changes. Further research should be conducted to investigate these relationships and their impact on the health and susceptibility to pathogens of small mammals, as they are significant reservoirs of zoonotic agents and frequently come into contact with humans in urban areas.

## Funding

This research was funded through the 2018-2019 BiodivERsA joint call for research proposals, under the BiodivERsA3 ERA-Net COFUND programme, and with the funding organization ANR.

## CRediT authorship contribution statement

MB: Conceptualization; Investigation; Data curation; Formal analysis; Methodology; Visualization; Writing - original draft

MG: Conceptualization; Methodology; Investigation; Supervision; Data curation; Formal analysis; Writing - review & editing

JP: Data curation; Resources

JF: Formal analysis; Resources

RG: Formal analysis; Writing - review & editing

AL: Formal analysis

BR: Conceptualization; Funding acquisition; Methodology; Supervision; Validation; Writing - review & editing

NC: Project administration; Conceptualization; Funding acquisition; Methodology; Supervision; Validation; Writing - review & editing

## Declaration of competing interest

The authors declare that they have no known competing financial interests or personal relationships that could have influenced the work reported in this paper.

## Appendix A. Supplementary data

Supplementary material

## Data availability

Raw data and scripts are available on zenodo repository (https://zenodo.org/record/8143272)

## Acknowledgments

We are grateful to all people that helped for field assistance, and to S. Piry for his help with data curation. We also thank J.F. Martin for his help with bioinformatics and microbiota analyses. Data used in this work were partly produced at the GENSEQ platform through the genotyping and sequencing facilities of ISEM (Institut des Sciences de l’Evolution-Montpellier) and Labex CeMEB (Centre Méditerranéen Environnement Biodiversité).

## Notes

https://zenodo.org/record/8143272

### Competing Interest Statement

The authors have declared no competing interest.

