## Supplementary Figure 1 for "Effects of forest urbanization on the interplay between small mammal communities and their gut microbiota"

**^c^** MIVEGEC, IRD, CNRS, Univ Montpellier, Montpellier, France

***Corresponding author at: Centre de Biologie pour la Gestion des Populations, 750 avenue agropolis, 34988 Montferrier sur Lez, France.**

Supplementary Figure 1

**Fig. S1.** Phylogeny of small mammals. Phylogenetic tree of small mammal species trapped in this study, built from *TimeTree of Life 5* (Kumar et al., 2022). The taxonomic ranks (order, family, genus and species) have been annotated on the tree. Nodes correspond to hypothetical common ancestors. Colors and silhouettes illustrate genus or families.

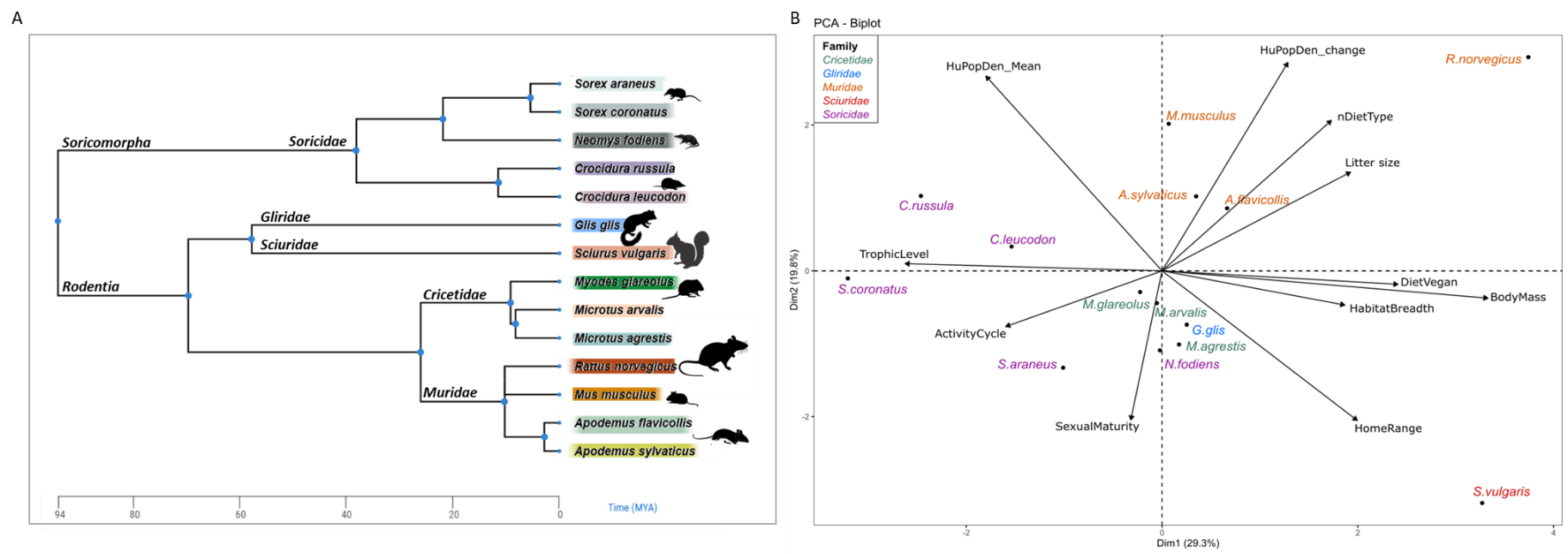
