## Supplementary Figure 2 for "Effects of forest urbanization on the interplay between small mammal communities and their gut microbiota"

**^c^** MIVEGEC, IRD, CNRS, Univ Montpellier, Montpellier, France

***Corresponding author at: Centre de Biologie pour la Gestion des Populations, 750 avenue agropolis, 34988 Montferrier sur Lez, France.**

Supplementary Figure 2

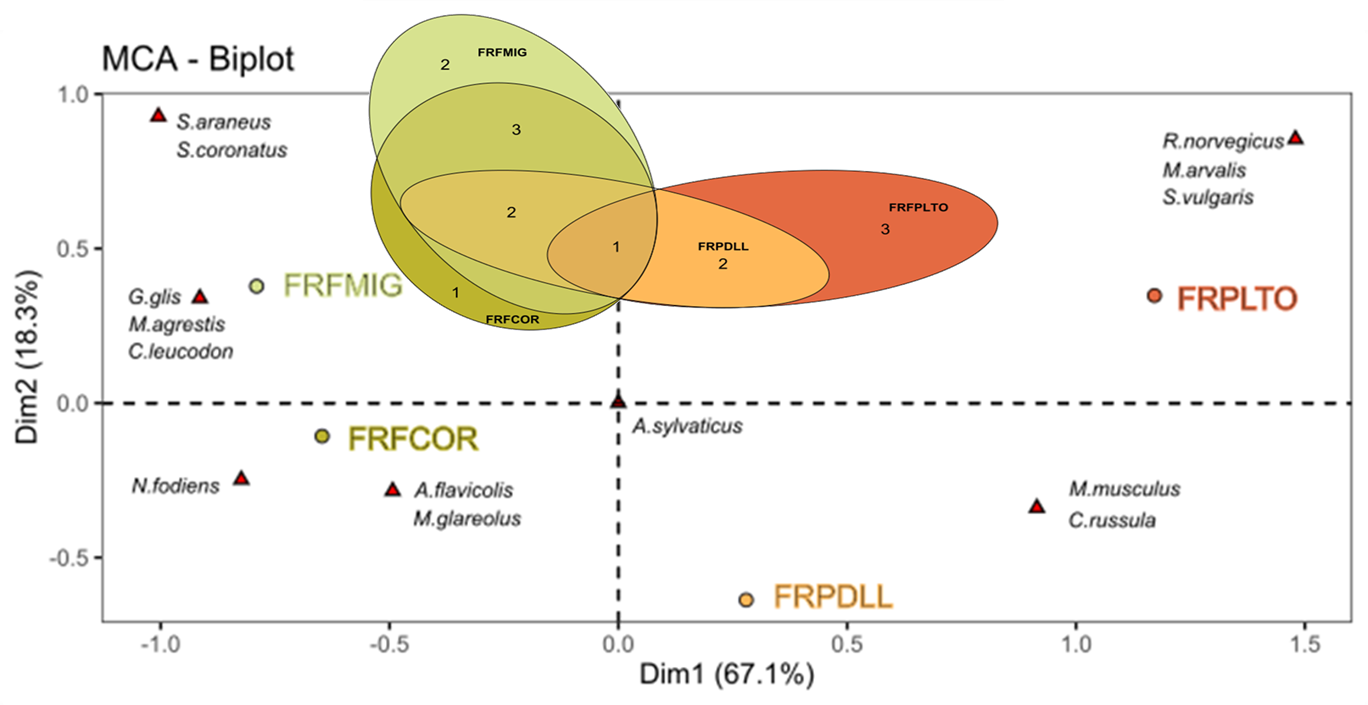

**Fig. S2**. Multiple correspondence analysis (MCA) of small mammal species present (represented by a red triangle) by sites (colored dots, FRFMIG and FRFCOR for the rural sites; FRPDLL and FRPLTO for urban sites). The Euler diagram shows more precisely the number of species shared between sites.
