## Supplementary Figure 3 for "Effects of forest urbanization on the interplay between small mammal communities and their gut microbiota"

**^c^** MIVEGEC, IRD, CNRS, Univ Montpellier, Montpellier, France

***Corresponding author at: Centre de Biologie pour la Gestion des Populations, 750 avenue agropolis, 34988 Montferrier sur Lez, France.**

Supplementary Figure 3

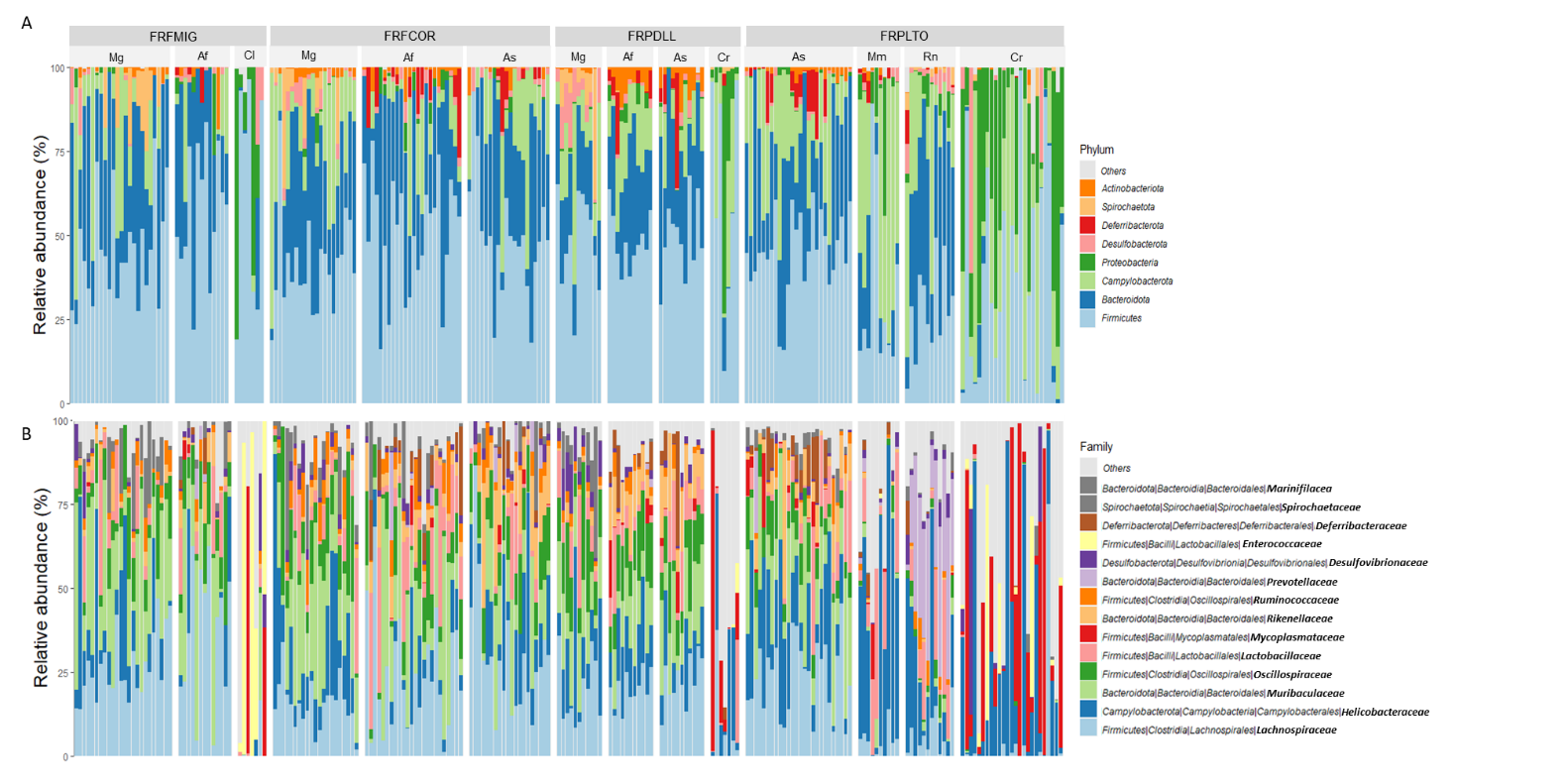

**Fig. S3.** Bar plot of the relative abundance (%) of A) Phylum and B) Family of small mammal gut microbiota, for each “species-site” combination. The Family legends correspond to Kingdom/Phylum/Class/Order/Family.
