## Supplementary Figure 4 for "Effects of forest urbanization on the interplay between small mammal communities and their gut microbiota"

**^c^** MIVEGEC, IRD, CNRS, Univ Montpellier, Montpellier, France

***Corresponding author at: Centre de Biologie pour la Gestion des Populations, 750 avenue agropolis, 34988 Montferrier sur Lez, France.**

Supplementary Figure 4

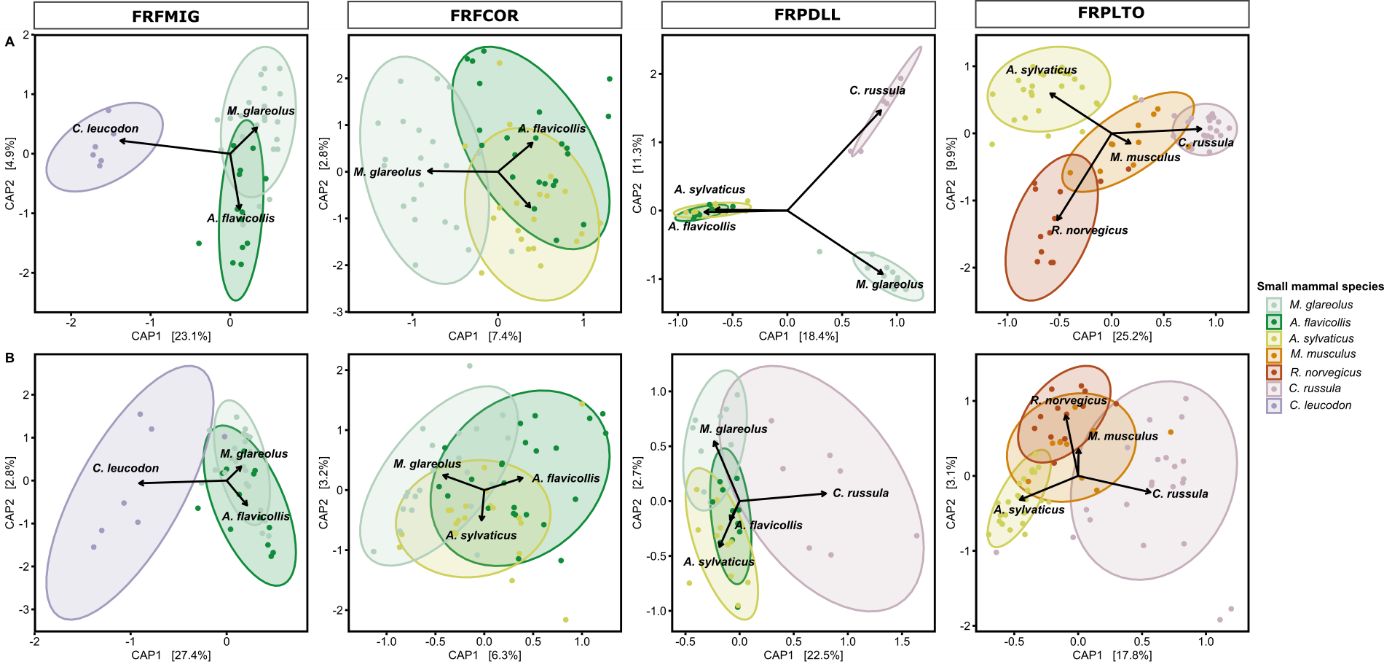

**Fig. S4.** Distance-based redundancy analysis (db-RDA) based on the composition of the gut microbiota of small mammal species for each sampling site, performed A) using ASVs and the weighted Unifrac dissimilarity matrix and B) using functions and the Bray-Curtis dissimilarity matrix. Only significant factors based on the *capscale* and *ordiR2step* analyses are indicated by arrows. The ellipses represent a 90% confidence interval around the centroids of the groups. Each point represents the gut microbiota of an individual and the color illustrates the small mammal species. Sites are ordered according to the urbanization gradient.
