## Supplementary Figure 5 for "Effects of forest urbanization on the interplay between small mammal communities and their gut microbiota"

**^c^** MIVEGEC, IRD, CNRS, Univ Montpellier, Montpellier, France

***Corresponding author at: Centre de Biologie pour la Gestion des Populations, 750 avenue agropolis, 34988 Montferrier sur Lez, France.**

Supplementary Figure 5

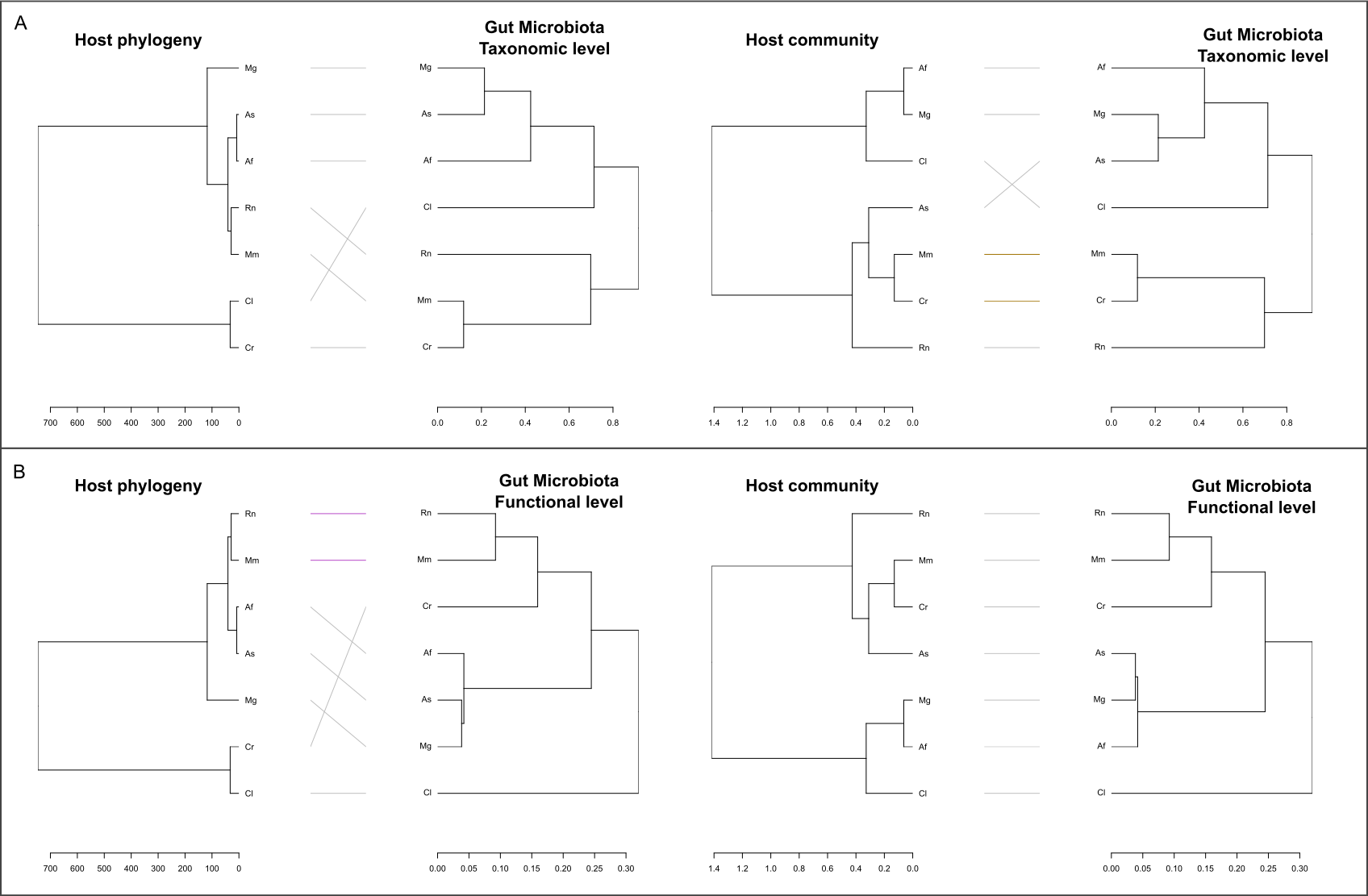

**Fig. S5** Tanglegrams showing concordance between 1) phylogenetic distance among small mammal and dissimilarities in function of their gut microbiota composition and 2) dissimilarities of niche for small mammal and dissimilarities in function of their gut microbiota composition. Concordance based on A) Taxonomy level (Weighted unifrac distance) and B) Functional level (bray curtis distance). The letters in tree correspond to Species ( Rn = R. *norvegicus*; Mm=*M. musculus*; As=*A. sylvaticus*; Af=*A. flavicollis*; Cr=*C. russula*; Cl=*C. leucodon*; Mg= *M. glareolus*)
