## Supplementary Figure 6 for "Effects of forest urbanization on the interplay between small mammal communities and their gut microbiota"

**^c^** MIVEGEC, IRD, CNRS, Univ Montpellier, Montpellier, France

***Corresponding author at: Centre de Biologie pour la Gestion des Populations, 750 avenue agropolis, 34988 Montferrier sur Lez, France.**

Supplementary Figure 6

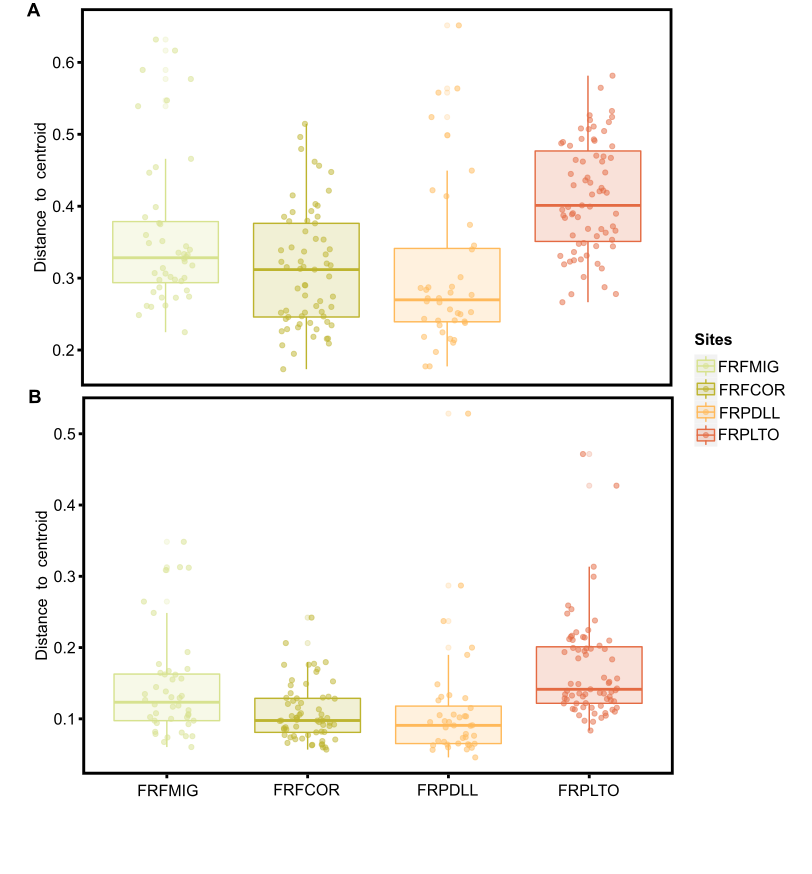

**Fig. S6.** Boxplot of the distance to the centroid of each individual within the group of sites at A) Taxonomic level and B) Functional level. Each point corresponds to an individual and the color to the sites (colored dots, FRFMIG and FRFCOR for the rural sites; FRPDLL and FRPLTO for the urban sites).
