## Supplementary Figure 7 for "Effects of forest urbanization on the interplay between small mammal communities and their gut microbiota"

**^c^** MIVEGEC, IRD, CNRS, Univ Montpellier, Montpellier, France

***Corresponding author at: Centre de Biologie pour la Gestion des Populations, 750 avenue agropolis, 34988 Montferrier sur Lez, France.**

Supplementary Figure 7

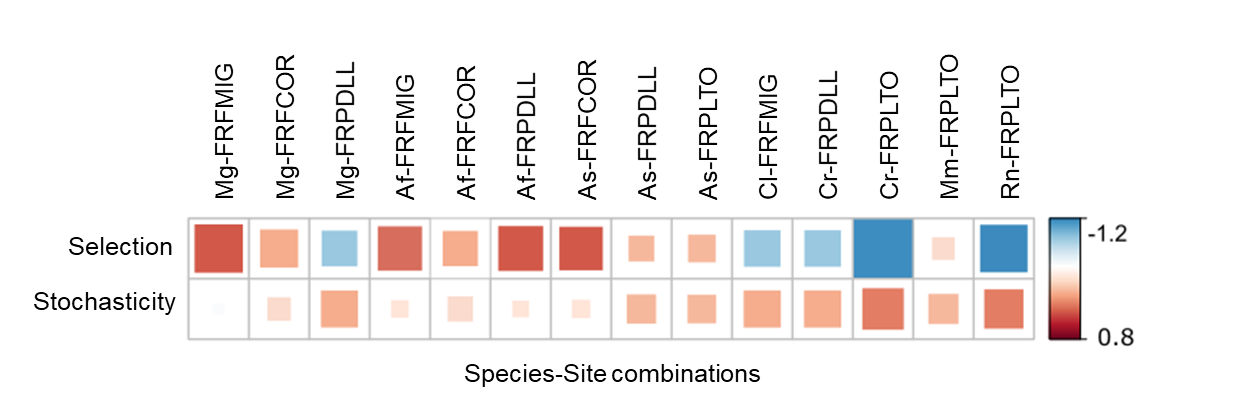

**Fig. S7.** Processes responsible for bacterial assembly and turnover in each combination of small mammal species and site. Corrplot representing the residual results of the chi-test analysis. The color gradient corresponds to the differences between observed and expected values. Negative values (blue) mean that processes are less observed than expected. Positive values (red) indicate that the share of processes is greater than expected.
