## Supplementary Figure 8 for "Effects of forest urbanization on the interplay between small mammal communities and their gut microbiota"

**^c^** MIVEGEC, IRD, CNRS, Univ Montpellier, Montpellier, France

***Corresponding author at: Centre de Biologie pour la Gestion des Populations, 750 avenue agropolis, 34988 Montferrier sur Lez, France.**

Supplementary Figure 8

**Fig. S8.** Heat map based on DESeq2 results illustrating significantly different abundances between species-site combinations, considering gut microbiota A) taxa (family level), B) metabolic pathways. The color gradient corresponds to the magnitude of the differences. Red values indicate a strong score and blue values a weak score. Species-site combinations and enzyme classes are ordered according to their profile.

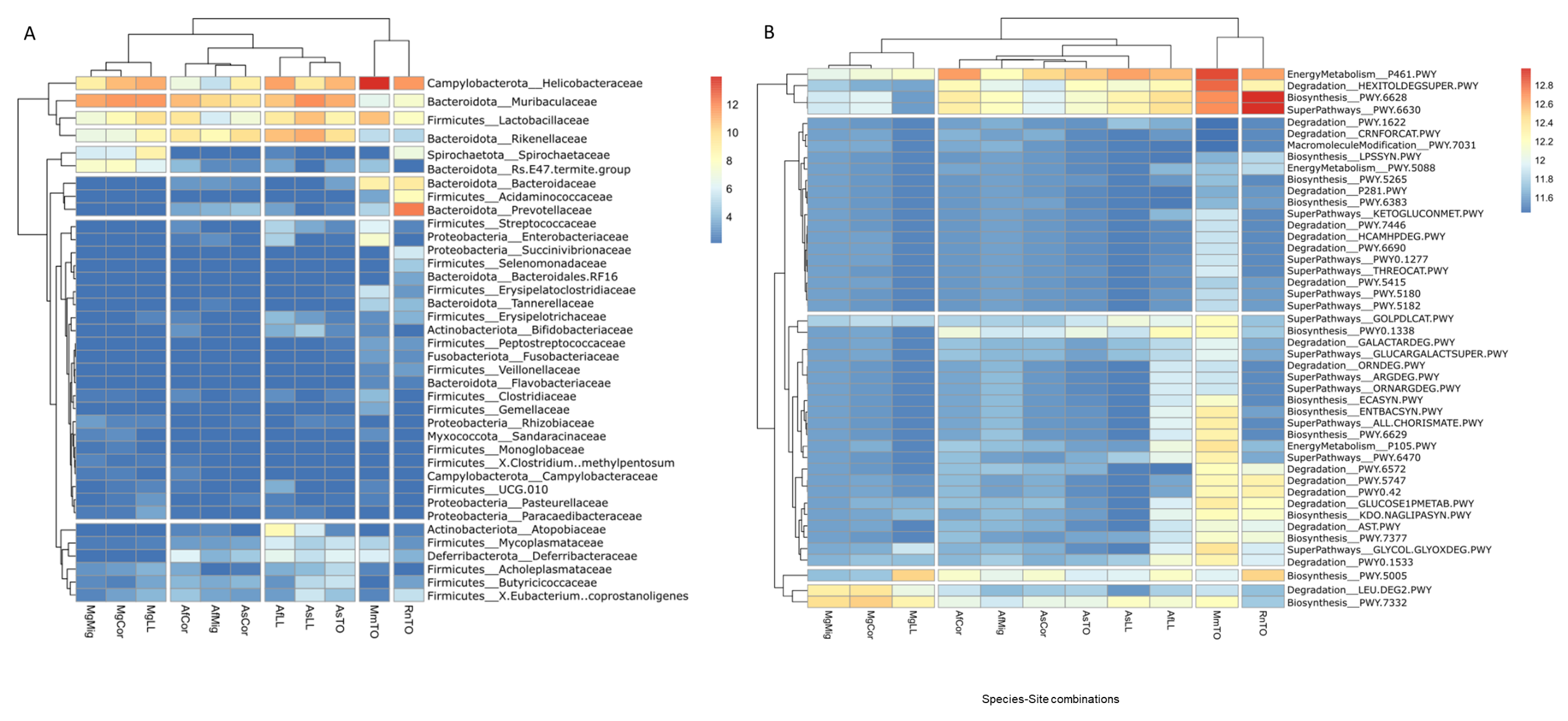
