## Supplementary File for "Effects of forest urbanization on the interplay between small mammal communities and their gut microbiota"

**^c^** MIVEGEC, IRD, CNRS, Univ Montpellier, Montpellier, France

***Corresponding author at: Centre de Biologie pour la Gestion des Populations, 750 avenue agropolis, 34988 Montferrier sur Lez, France.**

### Supplementary File

Description: Analyses of the differences between the gut microbiota of small mammals found dead or alive in traps

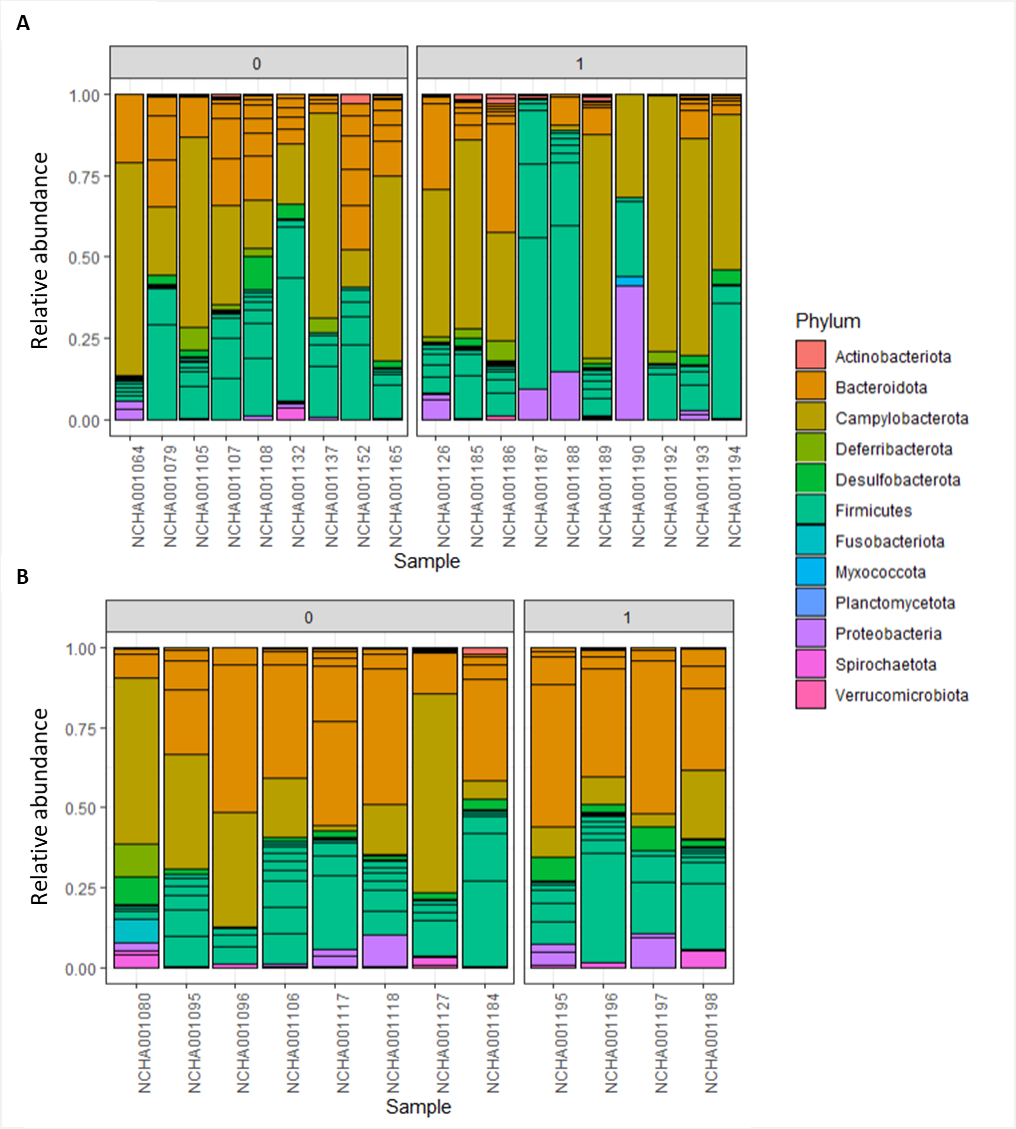

**Supp. File 1**. Differences in the relative abundance of ASVs at the phylum level between small mammals according to their status (code 0 = found alive in the trap, code 1= found dead in the trap) for A) *Mus musculus* and B) *Rattus norvegicus*. Each color corresponds to a phylum.

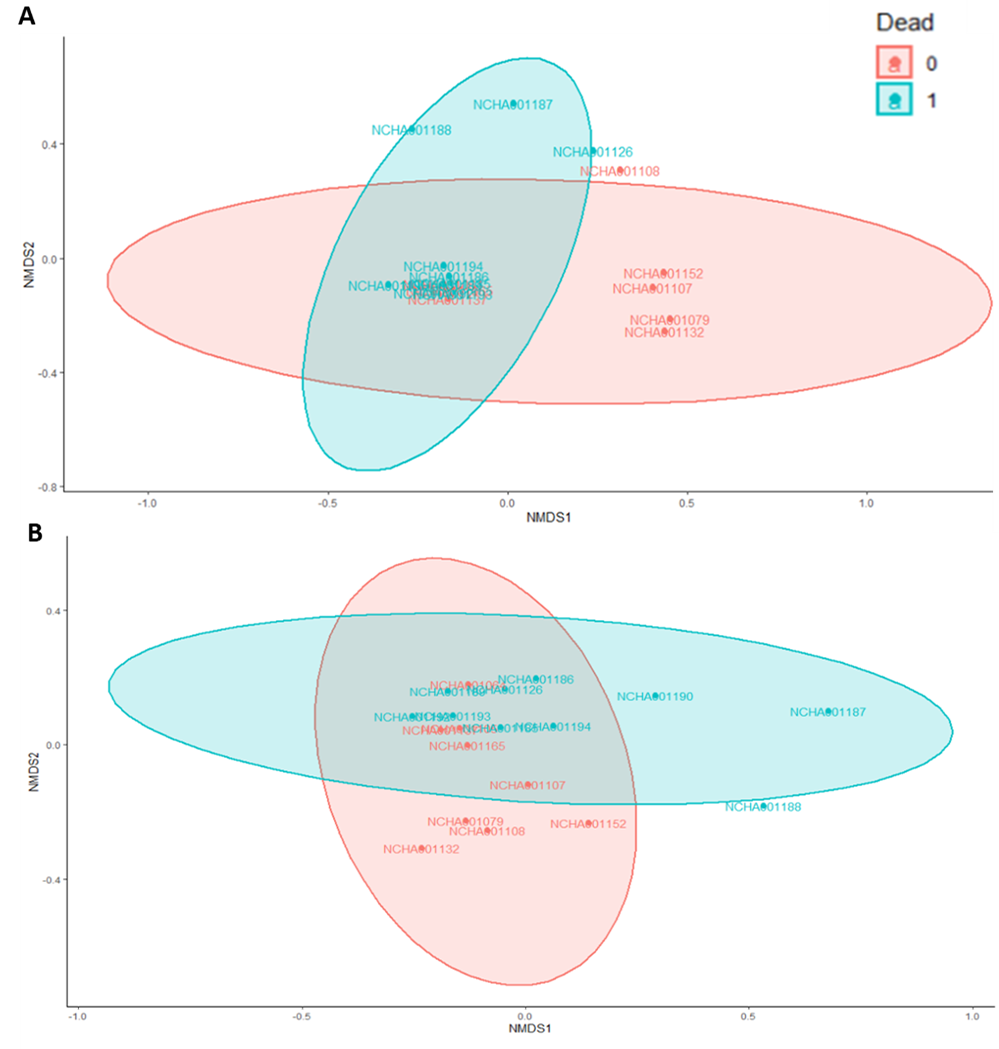

**Supp. File 2**. Differences in the composition of the GM (ASVs) at the phylum level according to small mammal status (code 0 = found alive in the trap, code 1= found dead in the trap) for A) *Mus musculus* and B) *Rattus norvegicus*

**Supp. File 3.** Statistical analysis (Adonis test) of the difference of GM composition between small mammals found dead or alive in the traps, for A) *Mus musculus* and B) *Rattus norvegicus*.

A) *Mus musculus*

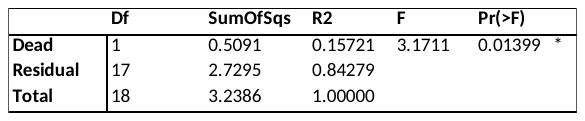

B) *Rattus norvegicus*

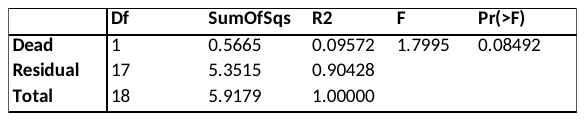

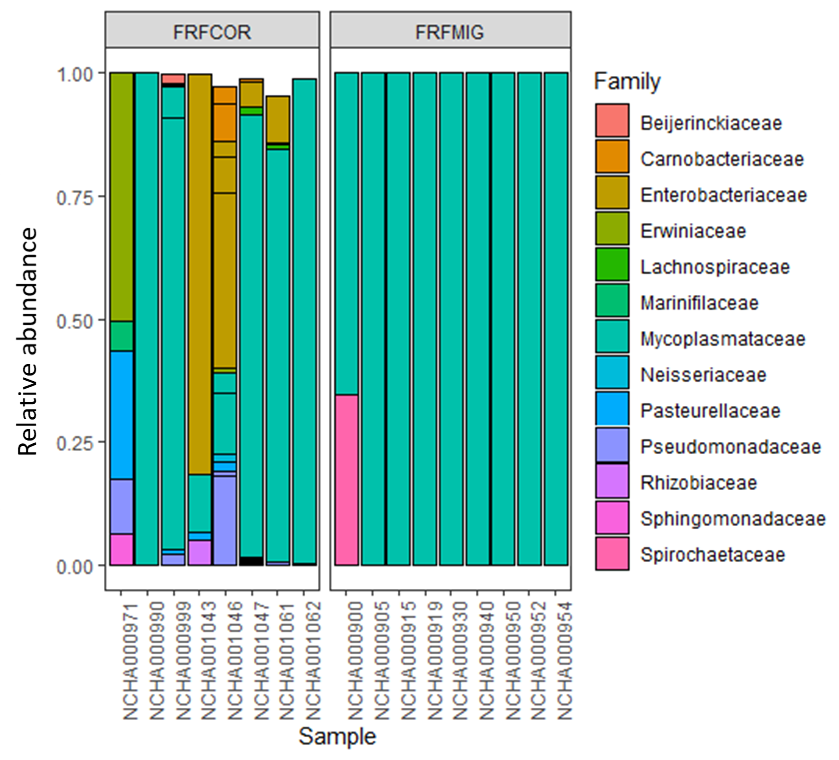

**Supp. File 4.** Relative abundance of GM taxa at the family level in *Glis glis* (each color corresponding to one family). We can observe an overabundance of the family *Mycoplasmataceae*, only few individuals had other taxa.
